# Serotonin signaling by maternal neurons upon stress ensures progeny survival

**DOI:** 10.1101/2020.01.20.913038

**Authors:** Srijit Das, Felicia K. Ooi, Johnny Cruz Corchado, Leah C. Fuller, Joshua A. Weiner, Veena Prahlad

## Abstract

Germ cells are vulnerable to stress. Therefore, how organisms protect their future progeny from damage in a fluctuating environment is a fundamental question in biology. We show that in *Caenorhabditis elegans*, serotonin released by maternal neurons during stress ensures the viability and stress tolerance of future offspring by enabling the transcription factor HSF1 to alter chromatin in soon-to-be fertilized germ cells by recruiting the histone chaperone FACT, displacing histones, and initiating protective gene expression. Without maternal serotonin signaling by neurons, FACT is not recruited by HSF1 in germ cells, transcription occurs but is delayed, and progeny of stressed *C. elegans* mothers fail to complete development. Serotonin acts through a signal transduction pathway conserved between *C. elegans* and mammalian cells to facilitate HSF1 to recruit FACT. These studies uncover a novel mechanism by which stress sensing by neurons is coupled to transcription response times of germ cells to protect future offspring.

## Introduction

The ability to react rapidly to environmental challenges is critical for the survival of individuals and species. In organisms with a nervous system, sensory neuronal circuits initiate many of the animals’ responses to environmental stressors, modifying behavior and physiology to adapt to the altered circumstance. However, whether and how sensory information used by the organism to predict impending danger is coupled to the protection of its future offspring is largely unknown. One conserved signaling molecule that is released in most organisms including *C. elegans*, early in response to real or perceived threats is the neuromodulator serotonin (5-hydroxytryptamine, 5-HT) ^1–7^. 5-HT is a bioamine secreted by specific neurons, and in some cases by peripheral cells, to modify learning and memory, behavior, development and physiological processes^1–8^, facilitating the animals’ future response to the stressor. For instance, in *Aplysia*, 5-HT increase mediates the encoding of memory required for habituation to a specific stressor and non-associative learning^9^. In mammals, increased 5-HT plays a dominant role in learning following social stress^7,10^. In *C. elegans*, we and others have shown that enhanced 5-HT mediates learned avoidance and activates defense responses. For example, pathogen odors increase 5-HT levels in *C. elegans* through the activity of chemosensory neurons^11–14^, and this increase in 5-HT is required for both the animal’s subsequent avoidance of pathogens, and its protection from infection. Similarly, exposure to increasing temperatures enhances 5-HT release from the serotonergic neurons (called NSM and ADF neurons) through the activity of the animal’s thermosensory neurons (called AFD neurons) and this release of 5-HT cell non-autonomously protects the animal from proteotoxicity ^2,15–17^. However, whether 5-HT released by the parent upon the detection of stress protects germ cells and future progeny from stress is not known. In fact, mammalian studies are suggestive of the opposite role for elevated levels of 5-HT that accompany chronic stress in the parent, and increased 5-HT is thought to contribute to behavioral and psychiatric disorders such as schizophrenia, depression, and autism in progeny through as yet poorly understood mechanisms ^18–21^—a unexpected effect given that stress-induced release of 5-HT by neurons and other 5-HT synthesizing cells is a highly conserved phenomenon.

Here we asked whether the stress-induced release of 5-HT by maternal neurons provides any benefits to germ cells and the development of future progeny. We used *C. elegans* to address this question in an *in vivo* setting, and cultured mammalian cells to dissect the molecular pathways by which 5-HT might act and examine the extent to which 5-HT mediated effects are conserved. We show that in *C. elegans*, 5-HT released by maternal neurons upon stress allows the transcription factor heat shock factor 1 (HSF1) to shorten the time to onset of mRNA production in soon-to-be fertilized germ cells. Specifically, 5-HT promotes the post-translational modification of HSF1 by protein kinase A (PKA) allowing HSF1 to recruit the histone chaperone FACT (<u>FA</u>cilitates <u>C</u>hromatin <u>T</u>ranscription) and alter histone dynamics to initiate transcription. This timely activation of HSF1 in germ cells ensures their viability and future stress tolerance: embryos that arise from heat-shocked mothers contain an excess of protective mRNA and are more tolerant to subsequent temperature insults as larvae. In the absence of maternal 5-HT, HSF1 activation in the germline is delayed, occurs without the recruitment of FACT, and a large fraction of embryos derived from these germ cells do not complete development, nor do they exhibit transgenerational thermotolerance. Remarkably, the intracellular signal transduction pathway by which 5-HT activates HSF1 enabling it to recruit FACT is conserved between *C. elegans* and mammalian cells. These results provide a novel mechanism by which 5-HT signaling protects germ cells, and developmental integrity. In addition they elucidate a molecular mechanism by which transcription response times of specific cells in a metazoan are tuned to stimulus intensity and onset.

## Results

### Maternal serotonin protects the germline from the detrimental effects of temperature stress

In *C. elegans* the only source of 5-HT is neuronal^22^. Tryptophan hydroxylase, TPH-1, the rate limiting enzyme for 5-HT synthesis, is expressed only in serotonergic neurons of hermaphrodites, and 5-HT synthesized and released by these neurons not only modifies neural circuit activity but is also distributed throughout the animal *via* the coelomic fluid to act on 5-HT receptors expressed by peripheral tissues ^23–25^. A deletion mutant in *tph-1* is viable and grossly wild-type, although completely devoid of 5-HT and therefore deficient in all responses that require 5-HT^22^. Therefore, to examine whether 5-HT released by maternal neurons upon the sensing of stress affected germ cells, we exposed wild-type animals and *tph-1* mutant animals to a transient and brief temperature-stress that we had previously shown enhances 5-HT release (5 minutes exposure to 34°C; a temperature gradient of 1 °C ± 0.2 °C increase per minute; see Materials and Methods), and then, evaluated the survival and development of embryos that arose from the heat-shocked wild-type and *tph-1* mutant animals (Figure S1A). Examining the numbers of already-fertilized embryos, the partially cellularized oocytes in adult animals that were soon-to-be fertilized, and the numbers of embryos laid during, and following heat exposure allowed us to extrapolate that embryos laid 0-2 hours following heat treatment were the already-fertilized embryos present *in utero* when the mothers were subjected to heat shock, and those generated between 2-4 hours following maternal heat-shock would be generated from partially cellularized germ cell nuclei (oocytes) present in the syncytial germline during the transient heat shock (Figure S1 B-D). This was true for both wild-type and *tph-1* mutant animals, although consistent with previously published data that 5-HT is required to modulate egg laying rates^23^, control *tph-1* mutants laid variable numbers of eggs during a 2 hour interval (Figure S1 C, D).

In the absence of stress all the embryos laid by wild-type as well as *tph-1* mutant animals hatched and developed into gravid adults indicating that 5-HT is not required for the survival of embryos under normal growth conditions (Figure S1 E, F and Figure 1A). Wild-type and *tph-*1 gravid adults themselves also survived the 5 minute heat exposure with no visible signs of damage (n=46 experiments, 4-5 animals/experiment; % survival wild-type and *tph-1* mutant animals=100), consistent with previous reports^1,26–28^ that exposure to high temperatures (35°C-37°C) for longer durations (hours) is required to impact the survival of adult *C. elegans*. However, the brief exposure to temperatures of 34°C was enough to disrupt embryonic development and ∼50% of embryos failed to hatch (Figure S1E, F). This was the case for embryos from wild-type or *tph-1* mutant mothers, irrespective of whether they were present *in utero* when the parents were subjected to the 5 minute-heat shock, or whether the embryos were first laid and then subjected to a 5-minute heat-shock (Figure S1E, F). Thus, it appeared that development processes were extraordinarily vulnerable to heat-induced disruption.

**Figure 1:**
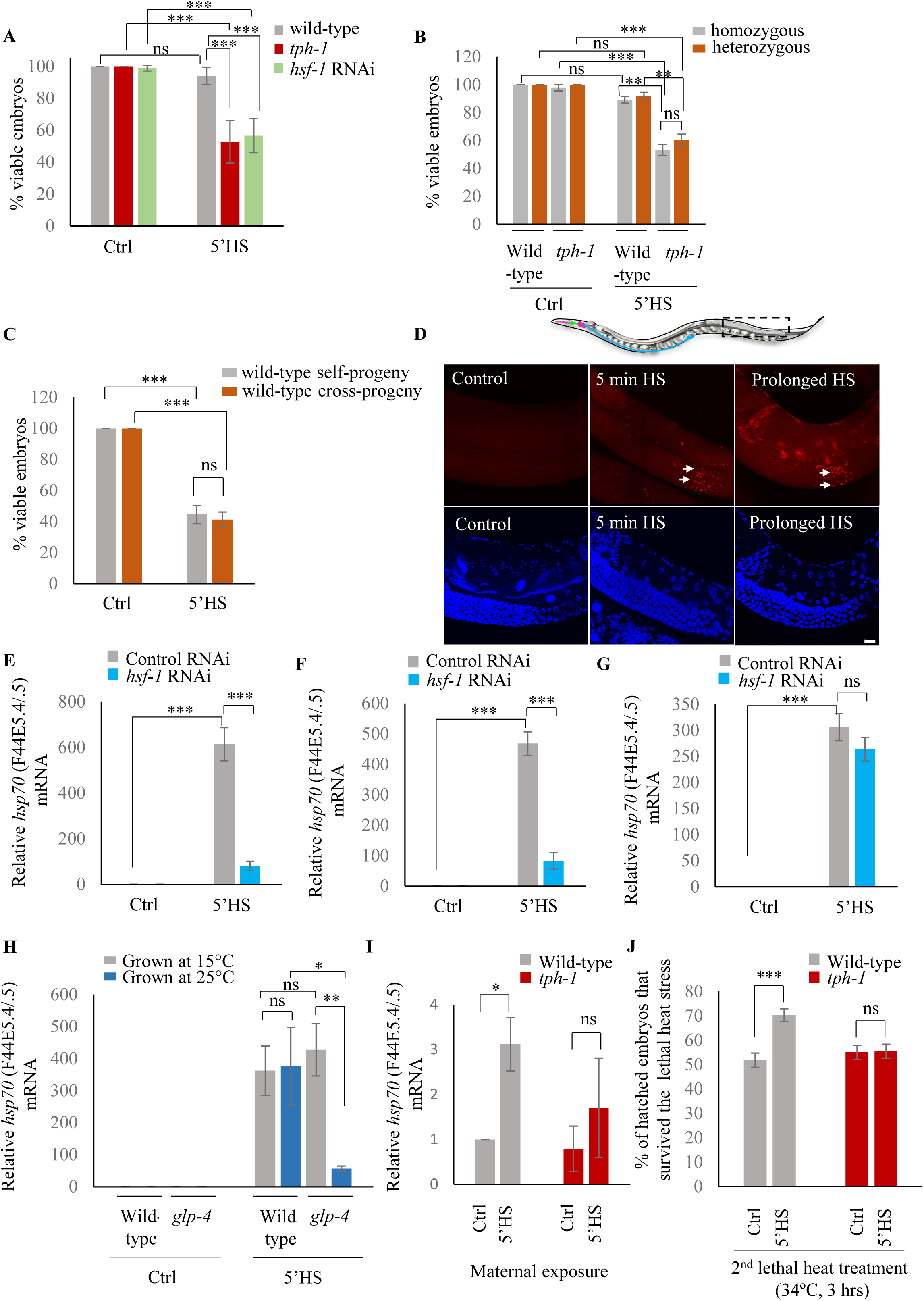
Maternal serotonin protects future progeny by enabling protective gene expression in the germline. **A**, Percent viable embryos from control (Ctrl) and heat-shocked (5’HS), wild-type animals, *tph-1* mutant animals and animals subjected to *hsf-1* RNAi. Embryos were laid during a 2-hour interval by non-heat shocked animals, or animals that were heat-shocked for 5 minutes at 34°C and allowed to recover at 20°C for 2 hours. Wild-type animals (n=28 experiments, embryos from 4-5 animals/experiment), *tph-1* mutant animals (n=28 experiments, embryos from 4-5 animals/experiment) and *hsf-1* RNAi-treated animals (n=5 control and 15 heat-shock experiments, embryos from 4-5 animals/experiment). **B,** Percent viable homozygous or heterozygous embryos from control (Ctrl) and heat-shocked (5’HS), wild-type and *tph-1* mutant animals. Wild-type and *tph-1* mutant hermaphrodites were allowed to mate with wild-type males and once the hermaphrodites were laying cross-progeny, embryos laid during a 2 hour interval by non-heat-shocked animals, or animals that had recovered for 2 hours post-heat-shock for 5 minutes at 34°C were scored. n=4 experiments, embryos from 4-5 animals/experiment. **C,** Percent viable embryos that were either self- or cross-progeny, laid by control (Ctrl) wild-type animals, or wild-type animals heat-shocked for 5 minutes at 34°C (5’HS). Wild-type hermaphrodites were allowed to mate with wild-type males that had been heat-shocked for 5 minutes at 34°C, and once the hermaphrodites were laying cross-progeny, the hermaphrodites were heat-shocked for 5 minutes at 34°C, and viable embryos laid 0-2 hours post-heat shock were scored (n=5 experiments, embryos from 4-5 animals/experiment). **D,** Representative confocal image showing *hsp70* (F44E5.4/.5) mRNA localization using smFISH in wild-type animals under control conditions and upon 5-minute and 15-minute (prolonged HS) exposure to 34°C (n=3 experiments, 24 animals). Optical sections were projected on one plane. Top, red: *hsp70* (F44E5.4/.5) mRNA. Bottom, blue: DAPI staining nuclei. Arrows indicate *hsp* mRNA in germline cells and arrowhead, in intestinal cells. Scale bar=10µm. **E-G,** Average *hsp70* (F44E5.4/.5) mRNA levels in control (Ctrl) and heat-shocked (5’HS) animals subjected to tissue specific RNAi. mRNA levels were normalized to that in control non-heat shocked, wild-type animals. Heat Shock: 5 minutes at 34°C. **E,** *rrf-1 (pk1714)* animals subjected to control RNAi and *hsf-1* RNAi (n=4 experiments). **F,** *mkcSi13* [*sun-1p::rde-1::sun-1 3′UTR* + *unc-119*(+)] *II; rde*-*1*(*mkc36*) *V* animals subjected to control RNAi and *hsf-1* RNAi (n=3 experiments). **G,** *rde-1(ne219) V; kbIs7* subjected to control RNAi and *hsf-1* RNAi (n=3 experiments)**. H,** Average *hsp70* (F44E5.4/.5) mRNA levels in control (Cntrl) and heat-shocked (5’HS) wild-type and *glp-4 (bn2) I* animals raised at 15°C (permissive temperature for *glp-4*) or 25°C (restrictive temperature for *glp-4*). n=4 experiments. mRNA levels were normalized to that in control non-heat shocked animals of same genetic background, raised at the corresponding temperature. Heat Shock: 5 minutes at 34°C. **I,** Average *hsp70* (F44E5.4/.5) mRNA levels in embryos laid during a 2 hours interval by wild-type or *tph-1* mutant animals that were not-heat shocked (Ctrl), or heat-shocked for 5 minutes at 34°C and allowed to recover for 2 hours (5’HS). n=3 experiments, embryos laid from 50 animals/experiment. mRNA levels were normalized to that in control non-heat shocked, wild-type embryos. **J.** Percent larvae from non-heat shocked, or heat-shocked wild-type and *tph-1* mutant animals that survive a subsequent heat exposure to 34 °C. Maternal heat shock: 5 minutes at 34°C. Larval heat shock: 3-hours at 34 °C. Note: larval heat exposure was titrated to achieve ∼50% lethality amongst progeny of control, non-heat shocked animals. n=5 experiments; larvae derived from 4-5 adult animals/experiment were scored. **A-C** and **E-J**: Data show Mean ± Standard Error of the Mean. ***, *p*<0.001; \*\**p*< 0.01, * *p*< 0.05 (paired Student’s t-test). ns, non-significant.

While the already-fertilized embryos of wild-type and *tph-1* mutant animals were susceptible to heat-shock, this was not the case for embryos derived from the fertilization of partially cellularized germ cells that were fertilized following heat shock of the parents and laid between 2-4 post-heat shock. These embryos survived and developed into adults—but only if they were derived from wild-type animals, and not from *tph-1* mutant animals (Figure 1A). Thus, while almost all the embryos (94 ± 2 %; n=28 experiments, embryos from 4-5 animals/experiment) generated from germ cells resident in wild-type animals during heat shock, but fertilized subsequently, hatched to develop into gravid adults (Figure 1A), only approximately 50% of the embryos generated similarly by germ cells in *tph-1* animals hatched (53 ± 2%; n=28 experiments, embryos from 4-5 animals/experiment; Figure 1A). This was surprising given the transient nature of the heat exposure and the fact that the germ cells were being fertilized and laid 2 hours following heat shock of the parents, at normal growth temperatures. Exposure of *C. elegans* to exogenous 5-HT causes 5-HT to be taken up into the serotonergic neurons and subsequently released, to an extent mimicking endogenous 5-HT^29^, although the kinetics of uptake and amounts released are not known. We tested whether such a treatment could rescue the lethality of these *tph-1* mutant embryos fertilized post-heat shock. Indeed, exposure of *tph-1* mutant mothers to exogenous 5-HT for even only 2 hours (during the 5-minute heat-shock treatment and the 2 hours until egg laying) rescued, in a significant manner, the lethality caused by the 5-minute heat-shock. (Figure S1 G). These data suggested that the presence of 5-HT protected the soon-to-be-fertilized germ cells from transient temperature fluctuations.

To investigate how 5-HT protected germ cells we asked whether the source of 5-HT that rescued embryonic lethality was maternal or embryonic, and whether it affected sperm or the female germline. To distinguish between a maternal and potentially embryonic sources, we examined the fate of embryos that were heterozygous for the *tph-1* mutant allele and therefore capable of synthesizing 5-HT, when laid by stressed *tph-1* mutant mothers devoid of 5-HT. If maternal 5-HT was responsible for the protective effects, these heterozygous embryos should remain equally susceptible to heat stress, despite being able to synthesize 5-HT. This was the case and embryos heterozygous for the *tph-1* mutant allele, generated by mating wild-type males with *tph-1* mutant hermaphrodites were equally susceptible to the 5-minute heat-shock as *tph-1* homozygous embryos when laid by stressed *tph-1* mutant mothers (Figure 1B). Thus, it appeared that 5-HT required for the viability of the germ cells was of maternal origin.

Since embryos of wild-type animals were susceptible to heat if they were fertilized prior to the heat-shock, but were resistant if they were fertilized following heat-shock, we asked whether survival was conferred by some sperm-derived factors generated by the 5-minute heat exposure. To do this, we assessed whether the embryos derived from oocytes fertilized by sperm from heat-shocked males survived the 5-minute heat shock. We verified that the embryos being assessed were indeed cross-fertilized with the heat-shocked sperm, and not self-fertilized by non-heat shocked sperm, by determining the sex ratios of the embryos laid by these mated mothers (see Materials and Methods). Irrespective of whether the embryos were generated by oocytes fertilized by heat-shocked sperm or by oocytes fertilized by non-heat shocked sperm, ∼50% of the embryos did not hatch if they were present *in utero* as post-fertilized embryos when the mothers were subjected to the 5 minute heat-shock (Figure 1C).

These data, together, showed that embryonic development was easily disrupted by temperature fluctuations, and maternal 5-HT protected the soon-to-be fertilized germ cells form stress-induced disruption, ensuring their survival. Furthermore, these data indicated that the effects of maternal 5-HT occurred, directly or indirectly, on the partially cellularized female germ cells of the parent hermaphrodite.

### Activation of HSF-1 in the germline is required to protect soon-to-be fertilized germ cells from heat stress

In all cells and organisms, a conserved and essential transcriptional response, the so-called ‘heat shock response’ counteracts the detrimental effects of heat or other stressors through the activation of the stress-inducible transcription factor, ‘heat shock factor 1’ (HSF1)^30–33^. This transcriptional response of HSF-1 (the sole *C. elegans* HSF1) was essential for the protection of germ cells upon heat exposure as, similar to *tph-1* mutant animals, nearly half the embryos (43 ± 4 % n=15 experiments, embryos from 4-5 animals/experiment; Figure 1A) generated from germ cells resident in animals subjected to RNA interference (RNAi)-induced knock-down of HSF-1 did not hatch when these animals were subjected to 5 minutes of temperature stress. In contrast almost all (93.7 ± 1%, n=3 experiments; embryos from 4-5 animals/experiment; Figure 1A) laid by *hsf-1* downregulated animals hatched and grew into mature adults in the absence of heat-shock showing that it was the heat-induced activity of HSF-1, and not its basal role that was required to protect germ cells. In addition, the adults with downregulated *hsf-1* themselves survived the 5-minute heat shock with no visible defects (n=15 experiments, 4-5 animals/experiment).

In agreement with the requirement for HSF-1 in stress-induced protection of embryos, a 5-minute exposure to heat was sufficient to activate HSF-1 and 408 genes, enriched in Biological Processes that handle misfolded proteins, were differentially expressed at 0.01 FDR by RNA-seq analysis of whole wild-type animals (Supplementary Figure S2A-C; Supplementary File 1). All these genes were, either directly or indirectly, dependent on HSF-1^34^ as a mutant that lacked the trans-activation domain of HSF1, previously shown to be viable but incapable of eliciting heat-induced transcriptional changes, showed no changes in gene expression upon heat shock (0.01 FDR; Figure S2B; Supplementary File 2). The differentially expressed genes included the major stress-induced *hsp70* genes, *hsp70* (F44E5.4/.5) and *hsp70* (C12C8.1), as well as other molecular chaperone genes that are the main targets of HSF-1 ^33,35^ and are known to counteract heat-induced damage (Figure S2C; Supplementary File 1).

Although the HSF-1-dependent transcriptional response upon heat shock has been well studied in *C. elegans*, the tissue-specificity of HSF-1-dependent gene expression in this metazoan is unclear. Importantly, it is not known whether germline cells express protective HSF-1-targets to account for their protection. Therefore, to examine whether HSF-1 was indeed activated in the germline upon the 5 minutes of heat-shock, we localized *hsp70* (F44E5.4/.5) mRNA in the whole animals following a 5 minute heat-shock using small molecule fluorescent *in situ* hybridization (smFISH). A 5-minute exposure to heat stress induced *hsp70* (F44E5.4/.5) mRNA predominantly in germline cells in late meiotic prophase and in very few other cells of the animal (Figure. 1D; Supplementary Figure 2D). These data suggested the HSF-1 targets were indeed activated in germline cells following a 5-minute heat-shock. Moreover, the germline was also amongst the first tissues to induce *hsp70* (F44E5.4/.5) mRNA, and mRNA was visible in germ cells after 5 minutes of heat-shock by smFISH, whereas continued exposure to heat was required to detect *hsp70* (F44E5.4/.5) mRNA in other cells of the animal (Figure 1D, Supplementary Figure 2D).

To confirm that *hsp* genes were indeed expressed in the germline upon 5 minutes of heat-shock, we utilized two additional methods. First, we knocked-down *hsf-1* in germline tissue using two different strains known to largely restrict all feeding-induced RNAi-mediated gene knock-down to germline tissue: (*mkcSi13* [*sun-1p::rde-1::sun-1 3′UTR* + *unc-119*(+)] *II; rde-1*(*mkc36*) *V*)^36^and *rrf-1 (pk1417)* ^37^mutants. We then examined the levels of *hsp70 (F44E5.4/.5)* induced following a 5-minute heat shock. We used two independent mutant backgrounds because tissue-specific RNAi in *C. elegans*, especially in the *rrf-1* mutants, has been shown to be leaky^38^ and can also occur in intestinal and epithelial cells. Second, we used a stain that harbors a temperature sensitive mutation in *glp-4* and fails to develop a germline at the restrictive temperature of 25°C (but has a fully functional germline at 15°C) and examined its transcription response to a 5 minute exposure to heat, to assess the contribution of the germline to the total amount of *hsp70* mRNA produced. Decreasing the levels of HSF-1 in the germline using the tissue-specific RNAi strains led to a marked decrease in the levels of *hsp70* (F44E5.4/.5) mRNA following a 5-minute heat-shock (Figure 1E, F). In contrast, the knock-down of *hsf-1* in intestinal cells, using a related tissue-specific RNAi strain (*rde-1(ne219) V; kbIs7*), did not significantly change the levels of *hsp* mRNA induced upon the 5-minute heat stress (Figure 1G). Similarly, abolishing the presence of germline cells using *glp-4* mutant animals grown at the restrictive temperature of 25 °C dramatically decreased *hsp70* (F44E5.4/.5) levels following a 5-minute heat-shock when compared to *glp-4* mutant animals that possessed functional germlines grown at permissive temperatures (15 °C) (Figure 1H). Moreover, *glp-4* mutant animals that possessed functional germlines because they were grown at permissive temperatures (15 °C) induced similar levels of *hsp* mRNA as wild-type animals that were grown at 15 °C (Figure 1H), confirming that these results was not a mere consequence of the change in the cultivation temperature. These data, together, indicated that the majority of *hsp70* mRNA produced by wild-type animals following the 5-minute heat shock was produced by germline cells.

Examination of the mRNA content of embryos laid by wild-type animals 2-4 hours after they had undergone a 5 minute heat shock releveled that these embryos had increased levels of *hsp70* (F44E5.4/.5) mRNA compared to embryos from control parents, perhaps accounting for their ability to survive the detrimental effects of heat (Figure 1I). In addition, the larvae that arose from these embryos displayed transgenerational protection from prolonged heat stress (Figure 1J). Specifically, when larvae were subjected to a 3-hour exposure to 34 °C, a condition titrated to achieve ∼50% lethality amongst progeny of control, non-heat shocked animals, significantly more larvae survived if they were progeny of heat-shocked mothers than if they were offspring of animals grown at control conditions (Figure 1J).

### Serotonin links the stress stimulus to the onset of protective gene expression

Upon the same 5 minute exposure to heat, *tph-1* mutant animals differentially expressed only 17, instead of 408 genes as measured by RNA-seq (Figure 2A; Figure S3A-C; Supplementary File 3) accumulated less *hsp* mRNA as measured by qPCR, (Figure 2B, C) and retained similar transcriptional profiles as unstressed *tph-1* animals by Principal Component Analysis (PCA; Supplementary Figure. 3D), indicating that they only mildly, if at all, activated HSF-1 in response to the transient temperature change. In contrast to wild-type embryos, embryos from heat-shocked *tph-1* mutant mothers also did not contain more *hsp70* mRNA (Figure 1I), nor did the larvae display increased stress tolerance (Figure. 1J). However, although *tph-1* mutant animals were deficient in the activating a heat-shock response upon a 5-minute exposure to heat stress, they were not deficient in activating HSF-1 *per se*. When exposed to greater intensities of heat stress (15 minutes exposure to 34°C, instead of 5 minutes), they accumulated similar levels of *hsp* mRNA as wild-type animals (Figure 2D, E). However, 15-minutes of heat exposure was already sufficient to impact the viability of embryos generated by germ cells as only 9 ± 1.5 % of the embryos from *tph-1* animals and 33 ± 5 % of the embryos from wild type animals (n=11 experiments, embryos from 4-5 animals/experiment) generated during the 2-4 hour time period after mothers had experienced 34°C for 15 minutes survived. (Figure 2F). This was the case despite the accumulation of equivalent levels of *hsp* mRNA in both wild-type and *tph-1* mutant animals upon a 15-minute heat shock. This suggested that in both wild-type and *tph*-*1* animals the germ cells were extremely vulnerable to heat damage, and the earlier onset of the protective heat shock response in germ cells that occurred in wild-type animals upon 5-HT release could maximize germ cell viability.

**Figure 2:**
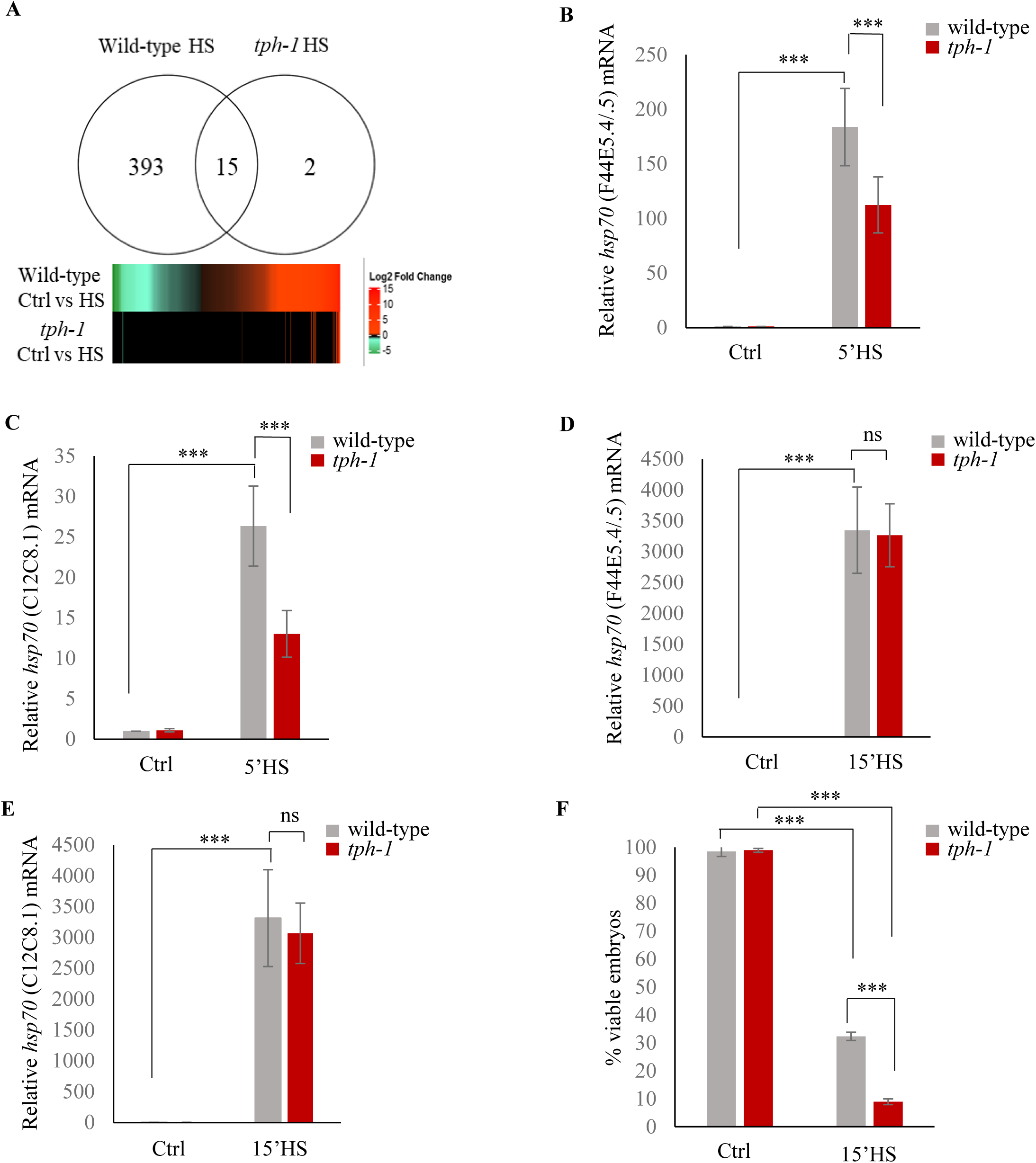
Serotonin accelerates the onset of gene expression upon heat shock. **A, top:** Venn diagram showing overlap between genes differentially expressed in wild-type animals and *tph-1* mutants (0.01 FDR) following 5 minutes heat-shock at 34°C. Numbers depict differentially induced genes in each strain. Data from RNA-seq experiments. **Bottom:** Heat map depicting expression levels (Log2 Fold change) of differentially expressed genes in wild-type and *tph-1* mutants. **B-E,** Average *hsp70* mRNA in wild-type and *tph-1* mutant animals following heat shock: **B,** *hsp70* (F44E5.4/.5) mRNA and **C,** *hsp70* (C12C8.1) mRNA levels in wild-type and *tph-1* mutant animals under control conditions and following heat shock at 34°C for 5 minutes (n=22 experiments). **D,** *hsp70* (F44E5.4/.5) mRNA and **E,** *hsp70* (C12C8.1) mRNA levels in wild-type and *tph-1* mutant animals under control conditions and following heat shock at 34°C for 15 minutes (n=5 experiments). **B-E**, mRNA levels were normalized to that in control non-heat shocked, wild-type animals. **F,** Percent viable embryos from control (Ctrl) and heat-shocked (15’HS), wild-type animals and *tph-1* mutant animals. Embryos were laid during a 2-hour interval by non-heat shocked animals, or animals that were heat-shocked for 15 minutes at 34°C and allowed to recover at 20°C for 2 hours. n=4 experiments, 4-5 animals/experiment. Data in **B-F** show Mean ± Standard Error of the Mean. ***, *p*<0.001 (paired Student’s t-test). ns, non-significant.

We have previously shown that in *C. elegans* 5-HT release acts cell-non autonomously to increase *hsp* gene expression. However, how 5-HT released by neurons activates transcription in remote tissues is not known. Therefore to ask how 5-HT may be modulating *hsp* expression in the germline, we used Chromatin immunoprecipitation (ChIP) followed by quantitative PCR (ChIP-qPCR)^39,40^ to assess the binding of key proteins involved in *hsp* transcription at *hsp* loci, in the presence and absence of 5-HT. Although we conducted ChIP in whole animals, we leveraged the fact that the majority of *hsp* transcription during 5 minutes of heat-shock occurred in the germ cells to infer that any changes in ChIP occupancy at *hsp* genes, if not reporting exclusively on what occurred in germline chromatin, would at the very least, be representative of changes at *hsp* loci in the germline.

The differences in the onset of transcription between wild-type animals and *tph-1* mutant animals was reflected by differences in the occupancy of RNA polymerase II (RNAP) and HSF-1 at *hsp70 (*F44E5.4/.5) and *hsp70* (C12C8.1), as assessed by ChIP-qPCR^39,40^ using primers sets targeted to the Promoter region of these *hsp* genes and, for RNAP, also to two regions within the gene body (Figure 3; Supplementary Figure 4A). In contrast to *Drosophila* and mammalian cells^41,42^, in *C. elegans*, RNAP pausing is rare in the absence of stress such as starvation^43,44^, and that was evident in the even distribution of RNAP across three distinct regions (‘Region A/Promoter, Region B and Region C) of the *hsp70* genes under basal, non-heat shock conditions (Supplementary Fig. 4A). In wild-type animals, consistent with the rapid induction of mRNA, RNAP was recruited to *hsp70 (*F44E5.4/.5) and *hsp70* (C12C8.1), peaking within 5 minutes of exposure to heat and remains significantly enriched at these genes upon continued heat exposure (Figure. 3A, C). The exact pattern of enrichment differed between the two *hsp70* genes, *hsp70 (*F44E5.4/.5) and *hsp70* (C12C8.1) promoters for reasons that are unclear. Notwithstanding these differences, in *tph-1* mutants RNAP occupancy was significantly lower than that in wild-type animals upon 5 minutes of heat shock (Figure 3A, C), but accumulated to similar or even higher levels as in wild-type animals upon continued heat exposure (15 minutes exposure to 34°C; Figure. 3A, C). Similarly, in wild-type animals HSF-1 was enriched at the promoter regions of *hsp* genes by 5 minutes of exposure of animals to heat, and remained enriched, although to lesser amounts by 15 minutes. In *tph-1* mutant animals HSF-1 was not recruited to these *hsp* gene promoters by 5 minutes, and HSF-1 levels at *hsp* promoters of *tph-1* mutants exposed to a 5 minute heat-shock did not significantly differ from that in control non-heat shocked animals. However, by 15 minutes following heat-shock, HSF-1 enrichment at *hsp* promoters in *tph-1* mutant animals was similar to that in wild-type animals after a 5-minute heat shock, suggesting that the binding of HSF-1 to its promoter was delayed in the absence of 5-HT (Figure 3B, D). The latter was evaluated by ChIP-qPCR using animals that expressed endogenous HSF-1 tagged at the C-terminus with 3X FLAG (Supplementary Fig. 4B). The enrichment of HSF-1 at target genes was specific and not apparent at the *syp-1* promoter that did not contain an HSF-1 binding site (Supplementary Fig. 4C), and was not a consequence of differences in HSF-1 protein levels between wild-type and *tph-1* mutants (Supplementary Fig. 4D, E).

**Figure 3:**
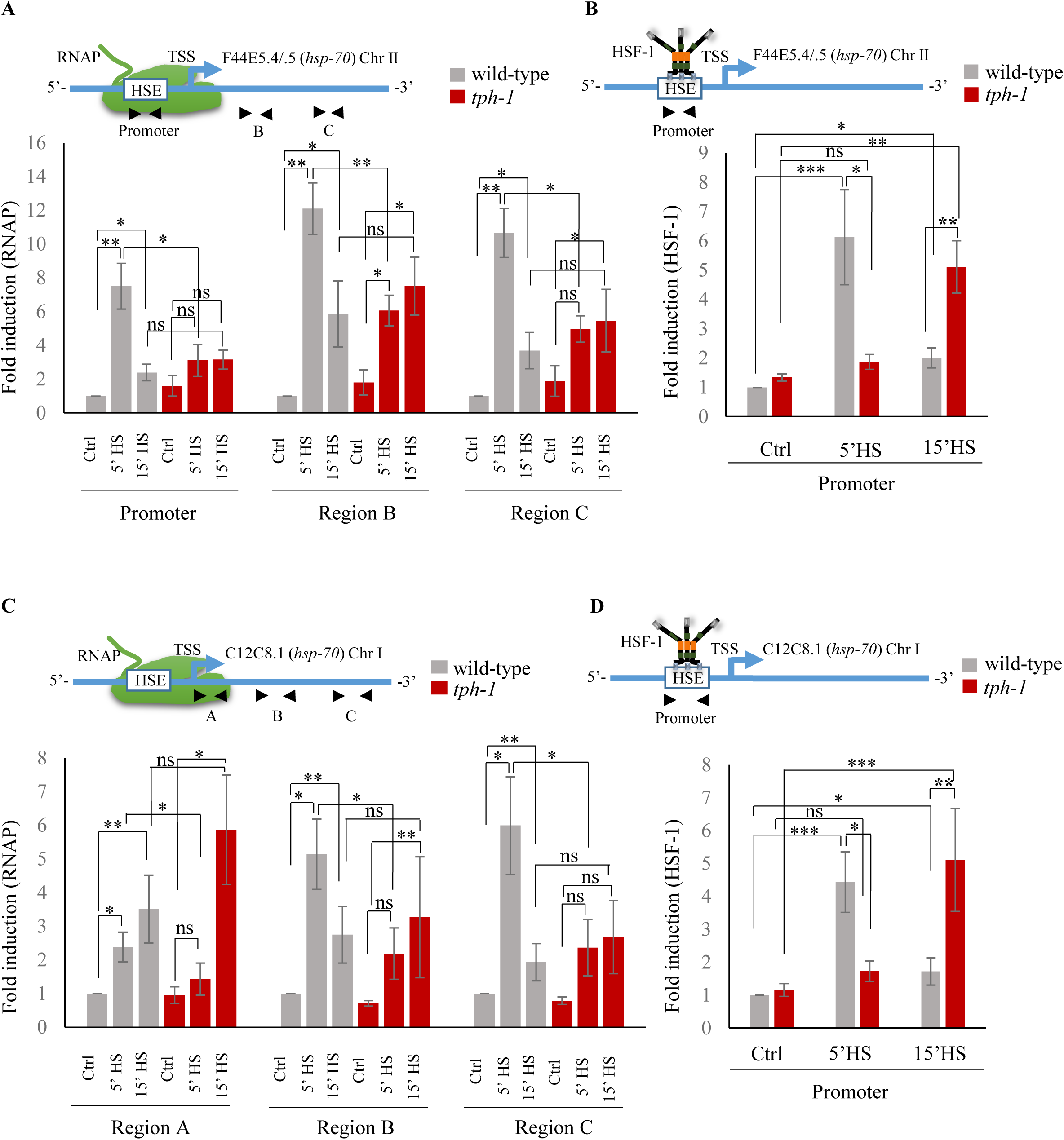
Serotonin accelerates the onset of RNAP and HSF-1 recruitment to target genes. **A, Top:** Schematic of *hsp70* (F44E5.4/.5) gene regions within the Promoter (−390 to −241), middle of gene (Region B: +696 to +915) and towards 3’-UTR (Region C: +1827 to +1996) that were assayed for occupancy by RNAP. **Bottom:** RNAP occupancy at Promoter, Region B and Region C in wild-type animals and *tph-1* mutants following exposure to 34°C for 5 minutes and 15 minutes (n=8 experiments). **B, Top:** Schematic of *hsp70* (F44E5.4/.5) gene regions within the Promoter (−390 to −241) assayed for occupancy of HSF-1. This is the same Promoter region as in **A**. **Bottom:** HSF-1 occupancy at the Promoter in wild-type animals and *tph-1* mutants following exposure to 34°C for 5 minutes and 15 minutes (n=14 experiments). **C, Top:** Schematic of *hsp70* (C12C8.1) gene regions close to the beginning (Region A: +25 to +185), middle of gene (Region B: +475 to +583) and towards 3’-UTR (Region C:+1645 to +1835) assayed for occupancy by RNAP. **Bottom:** RNAP occupancy at Region A, Region B and Region C in wild-type animals and *tph-1* mutants following exposure to 34°C for 5 minutes and 15 minutes (n=8 experiments). **D, Top:** Schematic of *hsp70* (C12C8.1) gene region within the Promoter (−166 to −78) assayed for HSF-1 occupancy. **Bottom:** HSF-1 occupancy at the Promoter in wild-type animals and *tph-1* mutants following exposure to 34°C for 5 minutes and 15 minutes (n=14 experiments). Data show Mean ± Standard Error of the Mean. Data in all experiments are normalized to values from control (non-heat shocked) wild-type animals. Specificity and efficiency of pull-down under control conditions was ascertained (see **Supplementary Figure 9**). *, *p*<0.05; **, *p*< 0.01 ***, *p*<0.001; (ANOVA with Tukey’s correction). ns, non-significant.

These data, together, supported a model whereby the release of maternal 5-HT in wild-type animals triggered an earlier onset of transcription. This difference in timing of the onset of transcription, was reflected by differences in the timing of HSF-1 and RNAP occupancy at *hsp* genes: in wild-type animals a 5 minute exposure was sufficient to induce a robust occupancy of HSF-1 protein and RNAP, while in the absence of 5-HT, binding of both HSF-1 and RNAP did occur, but was delayed. Along with previous data that the 5-minute exposure to heat activated HSF-1 dependent gene expression predominantly in germ cells of wild-type animals, these data suggested that 5-HT was responsible for the timing of HSF-1 activation in germ cells.

### Serotonin-dependent recruitment of FACT to displace histones hastens the onset of transcription

One mechanism by which 5-HT might accelerate the onset of transcription would be to alter chromatin accessibility to allow the transcription factor and RNAP^28,45–49^ to bind chromatin. To test if this was the case, we conducted ChIP-qPCR to examine levels of the histone H3, a component of the core nucleosome, at the two *hsp70* genes in wild-type animals and *tph-1* mutants upon transient exposure to heat (Figure 4A; Supplementary Figure 5A). In the absence of heat shock, H3 levels at the promoter, Transcription Start Site (TSS) and gene body (Region B) of *hsp* genes^49^ were comparable between wild-type animals and *tph-1* mutants, although *tph-1* mutant animals have slightly higher, but not significantly higher, levels throughout. In wild-type animals the brief exposure to heat disrupted histone-DNA interactions across the entire *hsp* gene: H3 occupancy at the promoter, TSS and gene body decreased significantly upon the 5 minute heat exposure as would be required to allow HSF-1 and RNAP access to DNA (Figure 4A; Supplementary Figure 5A). In contrast, in *tph-1* mutant animals that lack 5-HT, H3 occupancy did not decrease upon heat shock but instead remained similar to that under basal, non-heat shock conditions (Figure 4A; Supplementary Figure 5A). This suggested that changes in chromatin accessibility could underlie the differences in the response times of wild-type animals and *tph-1* mutants.

**Figure 4:**
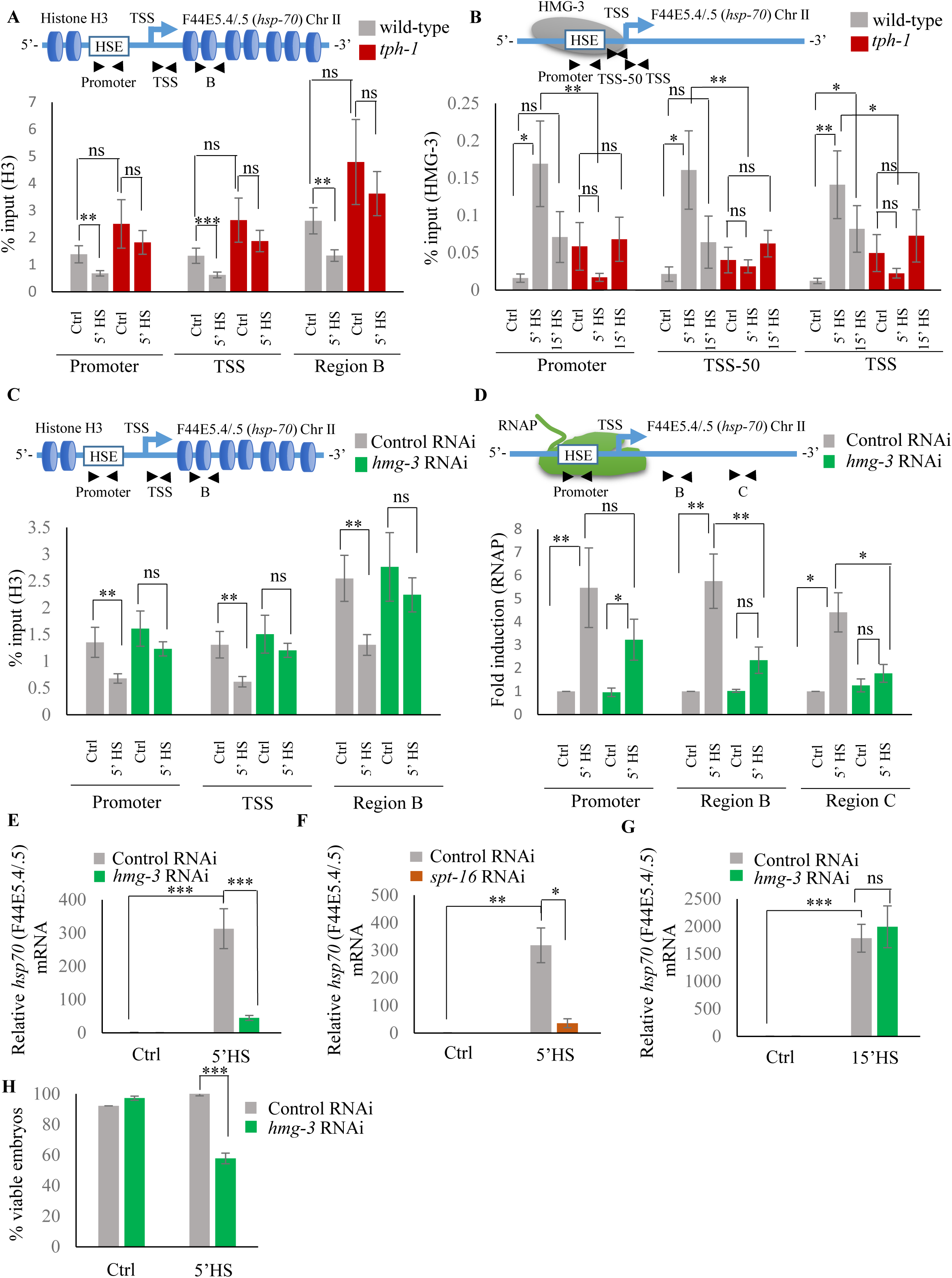
Serotonin enables FACT recruitment at *hsp* genes to displace nucleosomes and hasten the onset of transcription. **A, Top:** Schematic of *hsp70* (F44E5.4/.5) gene regions within the Promoter (same as Figure 3A and B: -390 to -241), Transcription Start Site (TSS: -81 to +38) and gene body (same as Region B in Figure 3A: +696 to +915) assayed for histone H3 occupancy. **Bottom:** Occupancy of histone H3 (% input) at Promoter, TSS and Region B upon 5 minutes at 34°C (n=9 experiments). **B, Top:** Schematic of *hsp70* (F44E5.4/.5) gene: regions in the Promoter (same as in Figure 3B: -390 to -241), region upstream of Transcription Start Site (TSS-50: -221 to -63) and Transcription Start Site (Same as **A**: TSS: -81 to +38) were assayed for HMG-3 occupancy. **Bottom:** HMG-3 occupancy (% input) across Promoter, TSS-50 and TSS following 5 and 15 minutes at 34°C (n=13 experiments). **C, Top:** Schematic of *hsp70* (F44E5.4/.5) gene regions assayed for H3 occupancy (same as in **A**). **Bottom:** Occupancy of histone H3 in Promoter, TSS and in Region B of *hsp70* (F44E5.4/.5) under control conditions and following 5 minutes at 34°C, in control-RNAi treated animals and *hmg-3-*RNAi treated animals (n=9 experiments). **D, Top:** Schematic of *hsp70* (F44E5.4/.5) gene: same regions as in Figure 3A, i.e. Promoter (−390 to −241), Region B (+696 to +915) and Region C (+1827 to +1996) were assessed for RNAP occupancy in control-RNAi treated and *hmg-3*-RNAi treated animals. **Bottom:** Fold change in RNAP across regions Promoter, Region B and Region C following 5-minute heat shock at 34°C (n=5 experiments). % input values were normalized to that in control-RNAi treated animals at Promoter and Regions B and C. Specificity and efficiency of pull-down under control conditions were verified (see **Supplementary Figure 9**). E, *hsp70* (F44E5.4/.5) mRNA levels in control-RNAi treated and *hmg-3* -RNAi treated animals following a 5-minute heat shock at 34°C (n=6 experiments). F, *hsp70* (F44E5.4/.5) mRNA levels in control-RNAi treated and *spt-16* -RNAi treated animals following a 5-minute heat shock at 34°C (n=4 experiments). G, *hsp70* (F44E5.4/.5) mRNA levels in control-RNAi treated and *hmg-3* -RNAi treated animals following a 15-minute heat shock at 34°C (n=6 experiments) **H,** Percent viable embryos (laid 2-4 hrs. post-5 minutes heat shock at 34°C) from control-RNAi treated and *hmg-3-*RNAi treated animals (n=5 experiments, 4-5 animals/experiment). Data show Mean ± Standard Error of the Mean. *, *p*<0.05; **, *p*< 0.01 ***, *p*<0.001; (**A-D**, ANOVA with Tukey’s correction; **E-H,** Paired Student’s t-test). ns, non-significant.

The histone chaperone FACT ^45,50–58^, a complex of two proteins, SPT16 and SSRP1, is known to disassemble histones to facilitate RNAP transcription at stress genes. In mammalian cells, FACT associates with HSF1 through its interaction with RPA (Replication Protein A), to allow the transcription factor access to DNA at the promoter^45^. In *C. elegans* the SSRP1 subunit of FACT consists of HMG-3 and HMG-4, of which HMG-3 is expressed exclusively in the germline, and HMG-4 in somatic tissue. HMG-3/HMG-4 along with SPT16, which is ubiquitously expressed, have been shown to displace nucleosomes and epigenetically modulate gene expression^59,60^. To investigate whether the observed difference in H3 loss between wild-type animals and *tph-1* mutants was mediated by FACT activity in germ cells, we examined HMG-3 occupancy at *hsp* genes in wild-type animals and *tph-1* mutant animals in strains expressing HMG-3 tagged at its endogenous locus with a 3X hemagglutinin (HA) tag^60^ (Figure 4B; Supplementary Figure 5B). Since HMG-3 is expressed exclusively in the germline of *C. elegans*, these data allowed us to make specific conclusions about the effects of 5-HT on germ cell chromatin.

As with HSF-1, HMG-3 protein levels are similar in wild-type and *tph-1* mutant animals (Supplementary Figure 5C, D). Nevertheless in wild-type animals, but not in *tph-1* mutants, HMG-3 was recruited to *hsp* genes by 5 minutes of heat shock (Figure 4B; Supplementary Figure 5B), and was necessary for the displacement of H3 histones at the Promoter, TSS and gene body as seen upon decreasing HMG-3 levels using RNAi (Figure 4C; Supplementary Figure 5E). HMG-3 was also necessary for RNAP occupancy at *hsp* genes upon 5 minutes of heat-exposure as RNAi-induced down-regulation of *hmg-3* decreased RNAP occupancy across the almost all regions of these genes. (Figure 4D; Supplementary Figure 5F). RNAP at Region A of *hsp70* (C12C8.1) was, for unknown reasons, not affected by *hmg-3* knock-down. Notwithstanding, the expression levels of *hsp* genes was diminished upon *hmg-3* RNAi (Figure 4E; Supplementary Figure 5G). A similar decrease in *hsp* gene expression upon the 5-minute heat shock was seen upon decreasing the levels of the HMG-3 interacting partner SPT-16 (Figure 4F; Supplementary Figure 5H), suggesting that HMG-3 and SPT-16 acted as a complex (FACT) to promote HSF-1 dependent gene expression in the germline.

In *tph-1* mutant animals HMG-3 was not recruited to *hsp* genes at significant levels either after 5 or 15 minutes of heat-shock (Fig. 4B; Supplementary Fig. 5B) suggesting that gene expression that occurred in *tph-1* mutants upon continued heat stress (Figure 2D, E) likely occurred through a HMG-3-independent mechanism. This was also supported by the observation that RNAi-induced downregulation of *hmg-3* levels in wild-type animals impaired *hsp* mRNA induction upon 5 minutes of heat shock (Figure 4E; Supplementary Figure 5G) but did not significantly affect *hsp* mRNA accumulation after 15 minutes (Figure 4G; Supplementary Figure 5I). Once again, even though HMG-3 was only required for the early onset of HSF-1 activation, and not for its activation *per se*, HMG-3 was required to protect germ cells from transient temperature fluctuations, as decreasing HMG-3 levels using RNAi decreased progeny survival upon transient heat shock much the same way as the lack of 5-HT or HSF-1 (Figure 4H).

The role of 5-HT in HMG-3 recruitment was confirmed by experiments where *hsp* gene expression^1,2^ was induced by optogenetically activating serotonergic neurons (ADF and NSM) to release 5-HT (Supplementary Figure. 6A-C). RNAi induced knock-down of *hmg-3* levels dramatically abrogated the 5-HT dependent increase in *hsp* mRNA (Figure. 5A; Supplementary Figure. 6D). In mammalian cells FACT is targeted to *hsp* promoters through its indirect interaction with HSF1 *via* RPA^45^. In *C. elegans* also FACT recruitment to *hsp* genes depended directly or indirectly on HSF-1 as RNAi-dependent downregulation of *hsf-1* decreased FACT recruitment at *hsp* genes (Figure 5B; Supplementary Figure. 6E). These data together indicated that 5-HT-signaling enabled HSF-1 to recruit HMG-3 in germ cells and displace histones to shorten the onset of RNAP-dependent gene expression and ensure viability of germ cells during stress.

**Figure 5:**
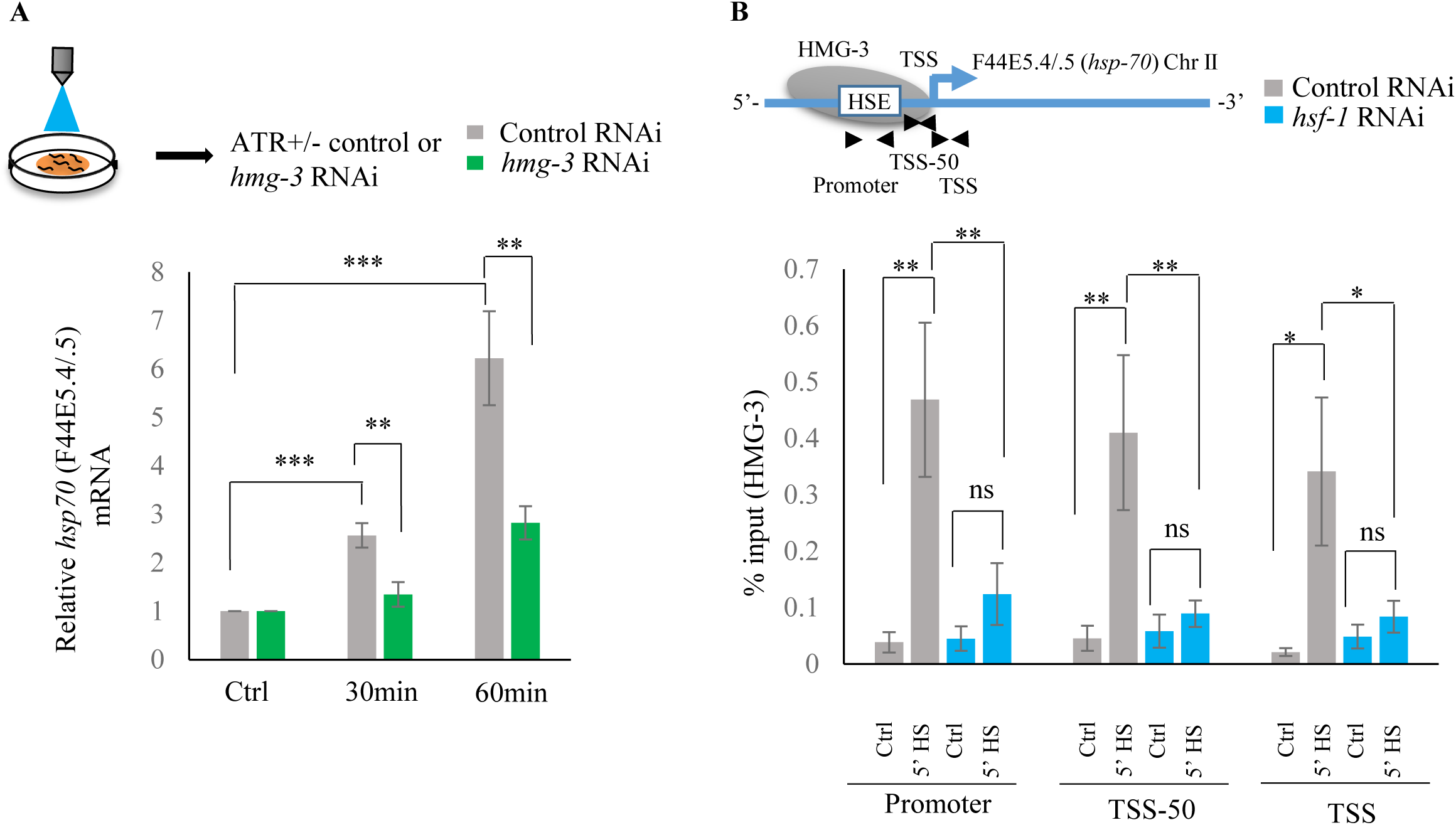
Serotonin release mediates FACT recruitment by HSF-1 to induce *hsp* expression. **A, Top:** Schematic of optogenetic activation of 5-HT release conducted by stimulating ADF and NSM neurons in control-RNAi treated and *hmg-3-*RNAi treated animals. **Bottom:** *hsp70* (F44E5.4/.5) mRNA levels in control-RNAi treated and *hmg-3-*RNAi treated animals at different time points following optogenetic stimulation. mRNA levels were normalized to control-RNAi treated and *hmg-3*-RNAi treated unstimulated animals respectively (n=6 experiments). **B, Top:** Schematic of *hsp70* (F44E5.4/.5) gene Promoter, TSS-50 and TSS to assess HMG-3 occupancy in control-RNAi treated and *hsf-1-*RNAi treated animals. **Bottom:** HMG-3 occupancy (% input) at Promoter, TSS-50 and TSS in control-RNAi and *hsf-1* - RNAi treated animals following 5 minutes at 34°C (n=9 experiments). Specificity and efficiency of pull-down under control conditions was ascertained (see **Supplementary Figure 9**). Data show Mean ± Standard Error of the Mean. *, *p*<0.05; **, *p*< 0.01 ***, *p*<0.001. **A**, Paired Student’s t-test. **B**, ANOVA with Tukey’s correction). ns, non-significant.

### Serotonin-induced PKA-activation is a conserved signaling pathway that enables HSF1 to recruit FACT

To identify the intracellular signal transduction pathway triggered by 5-HT to modulate the interaction of HSF-1 with FACT, we decided to use mammalian cells where we would be able to isolate cell autonomous effects away from cell non-autonomous effects, and leverage the wealth of information about mammalian HSF1 ^30,33, 61–66^ (Supplementary Figure 7A). As in *C. elegans*, exposure of mammalian cells to exogenous 5-HT could also autonomously activate HSF1. Treatment of mouse primary cortical neurons^67–69^ with exogenous 5-HT resulted in a dose- and time-dependent increase in mRNA levels of the most highly inducible *hsp* genes that are targets of mammalian HSF1: *Hspb1* (Supplementary Figure 7B), *Hspa1a* (Figure. 6A) and *Hspb5* (Supplementary Figure 7C). A similar increase in *HSPA1A* mRNA (Figure 6B) and HSPA1A protein levels (Supplementary Figure 7D, E) was observed upon treatment of human NTera2 (NT2) cells. siRNA induced knock-down of HSF1 (Supplementary Figure 7F) abrogated 5-HT-induced *HSPA1A* mRNA expression (Figure 6C). Thus, remarkably, acute increases in 5-HT activated HSF1-dependent gene expression in mammalian cells, much the same way it did in *C. elegans*.

**Figure 6:**
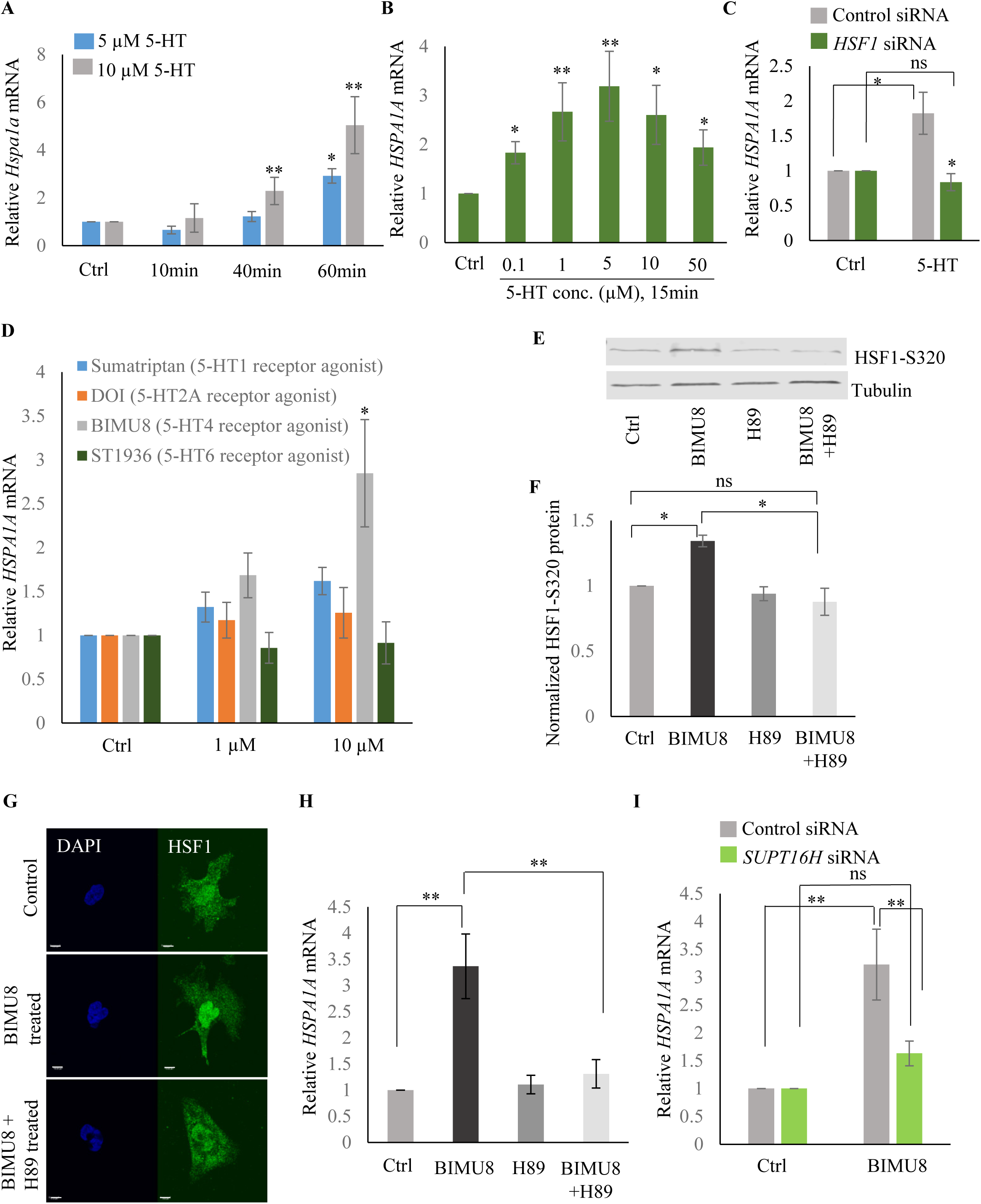
Serotonin activates PKA-mediated signal transduction to enable HSF1-FACT interaction in mammalian cells. **A,** Time and dose-dependent change in *Hspa1a* mRNA levels in control and 5-HT treated primary cortical neuronal cultures (n=4 experiments). **B,** Dose-dependent change in *HSPA1A* mRNA levels in control NT2 cells and NT2 cells treated with 5-HT for 15 minutes (n=4 experiments). **C,** *HSPA1A* mRNA levels in NT2 cells treated with 5µM 5-HT for 15 minutes, transfected with control and *HSF1* siRNA (n=4 experiments). **D,** *HSPA1A* mRNA levels in NT2 cells treated with two different doses of four 5-HT receptor agonists relative to control untreated cells (n=5 experiments). NT2 cells were treated for 10 minutes. **E-F,** Protein levels of S320 phospho-modified HSF1 in control NT2 cells and cells treated with 10µM BIMU8 for 10 minutes, in the presence or absence of the PKA inhibitor, H89 (n=4 experiments). **E,** Representative western blot using an antibody that recognizes HSF1 phosphorylated at S320. Tubulin served as the internal control. **F,** Quantitation of phospho-S320 levels (n=4 experiments). **G,** Representative micrographs showing projections of confocal images of HSF1 localization in control NT2 cells and cells treated with 10µM BIMU8 for 10 minutes, in the presence or absence of the H89 (n=2 experiments; 25 cells). Scale bar=10µm. **H,** *HSPA1A* mRNA levels relative to control NT2 cells upon treatment with 10µM BIMU8 for 10 minutes, in the presence or absence of H89 (n=5 experiments). **I,** *HSPA1A* mRNA levels in cells treated with 10µM BIMU8 for 10 minutes, transfected with control and *SUPT16H* siRNA. mRNA levels and protein levels are normalized to control RNAi-treated or unstimulated cells (n=5 experiments). Data in **A-D, F, H, I** show Mean ± Standard Error of the Mean. *, *p*<0.05; **, *p*< 0.01 ***, *p*<0.001; (Paired Student’s t-test). ns, non-significant.

The effects of 5-HT are transduced through intracellular signal transduction pathways and depend on the particular 5-HT receptor(s) involved in the biological process (either G protein-coupled receptors—GPCRs, or ligand-gated ion channels) ^70,71^. Therefore, to identify the intracellular pathway involved in 5-HT-induced HSF1 activation, we used a panel of 5-HT receptor specific agonists^10,16,25, 71–75^. Agonists of 5-HT4 receptors (BIMU8), but not 5-HT6, 5-HT2A and 5-HT1 elicited a dose- (Figure 6D) and time-(Supplementary Figure. 7G) dependent increase in *HSPA1A* mRNA in NT2 cells that was HSF-1 dependent (Supplementary Figure 7H), mimicking the effects of 5-HT. BIMU8 also induced *Hspa1a* and *Hspb1* mRNA in primary cortical neurons (Supplementary Figure 7I). The 5-HT4 receptor is a GPCR that signals though adenylyl cyclase and activates protein kinase A (PKA) ^73,74, 76–79^. PKA has been shown to phosphorylate mammalian HSF1 at the serine 320 residue during heat shock^80–82^. In agreement with this, BIMU8 treatment triggered an increase in S320-modified HSF1^81,82^ as detected by a phospho-specific antibody (Figure 6E, F). Inhibiting PKA activity using the drug H89^83^ inhibited the BIMU8-induced increase in S320 phosphorylation (Figure 6E, F). In addition, BIMU8 treatment promoted the nuclear localization of HSF1 which in turn could also be inhibited by H89, recapitulating previous studies on PKA-induced phosphorylation of HSF1^81^ (Figure. 6G; Supplementary Fig. 7J). *HSPA1A* mRNA levels that were induced by BIMU8 treatment were also inhibited upon treatment of the cells with H89 (Figure 6H). Moreover, as with 5-HT induced activation of HSF-1 in *C. elegans,* BIMU8-induced activation of HSF1 in NT2 cells also required FACT, and the knockdown of the *SUPT16H* subunit of FACT by siRNA (Supplementary Figure. 7K) abrogated the BIMU8-induced upregulation of *HSPA1A* mRNA (Figure 6I). These data together indicate that 5-HT cell-autonomously enables HSF1 to recruit FACT in mammalian cells through the activation of 5-HT4 receptor and the conserved cAMP-PKA intracellular signaling pathway, and as in *C. elegans* this allowed HSF1 to access *hsp* genes and initiate RNAP-dependent gene expression even in the absence of stress.

Although *C. elegans* do not possess a 5-HT4 receptor ortholog they possesses 5-HT receptors that activate PKA^23^. Therefore, to examine whether in *C. elegans* also, 5-HT acted through PKA to accelerate the onset of HSF-1-dependent gene expression we modulated the *C. elegans* PKA holoenzyme. PKA exists as a tetramer with catalytic and regulatory subunits, and the release of inhibition by the regulatory subunits results in the enabling of the catalytic activity of PKA. In *C. elegans kin-1* encodes the catalytic subunits of PKA, and inhibition of *kin-1* diminishes PKA activity, while *kin-2* encodes the regulatory subunits, and decreasing *kin-2* levels releases KIN-1 and activates PKA^84–89^. Decreasing *kin-1* levels by RNAi dampened the induction of *hsp70* mRNA that occurs upon optogenetic activation of 5-HT release (Figure 7A; Supplementary Figure. 8A). Decreasing *kin-1* levels by RNAi also prevented the recruitment of HMG-3 in germ cells by HSF-1 after transient exposure to heat (Figure 7B; Supplementary Figure 8B). Moreover, as in mammalian cells the role of KIN-1 in activating HSF-1 appeared to be cell autonomous, as decreasing *kin-1* levels only in germ cells decreased *hsp70* mRNA levels upon 5 minutes heat-shock, similar to decreasing *kin-1* levels in whole animals (Figure. 7C, D; Supplementary Figure 8C, D). Conversely, activating PKA in *tph-1* animals by knocking down *kin-2* rescued the delayed response of *tph-1* mutant animals, both increasing occupancy of HSF-1 at *hsp* promoters by 5 minutes upon heat exposure despite the absence of 5-HT (Figure 7E; Supplementary Fig. 8E), and increasing *hsp70* (F44E5.4/.5) mRNA levels to wild-type levels upon 5 minutes heat shock (Figure 7F; Supplementary Figure 8F). Activating PKA by RNAi mediated knockdown of *kin-2* also rescued, significantly albeit incompletely, the embryonic lethality induced by exposing *tph-1* mutant animals to 5 minutes of heat (Supplementary Figure. 8G).

**Figure 7:**
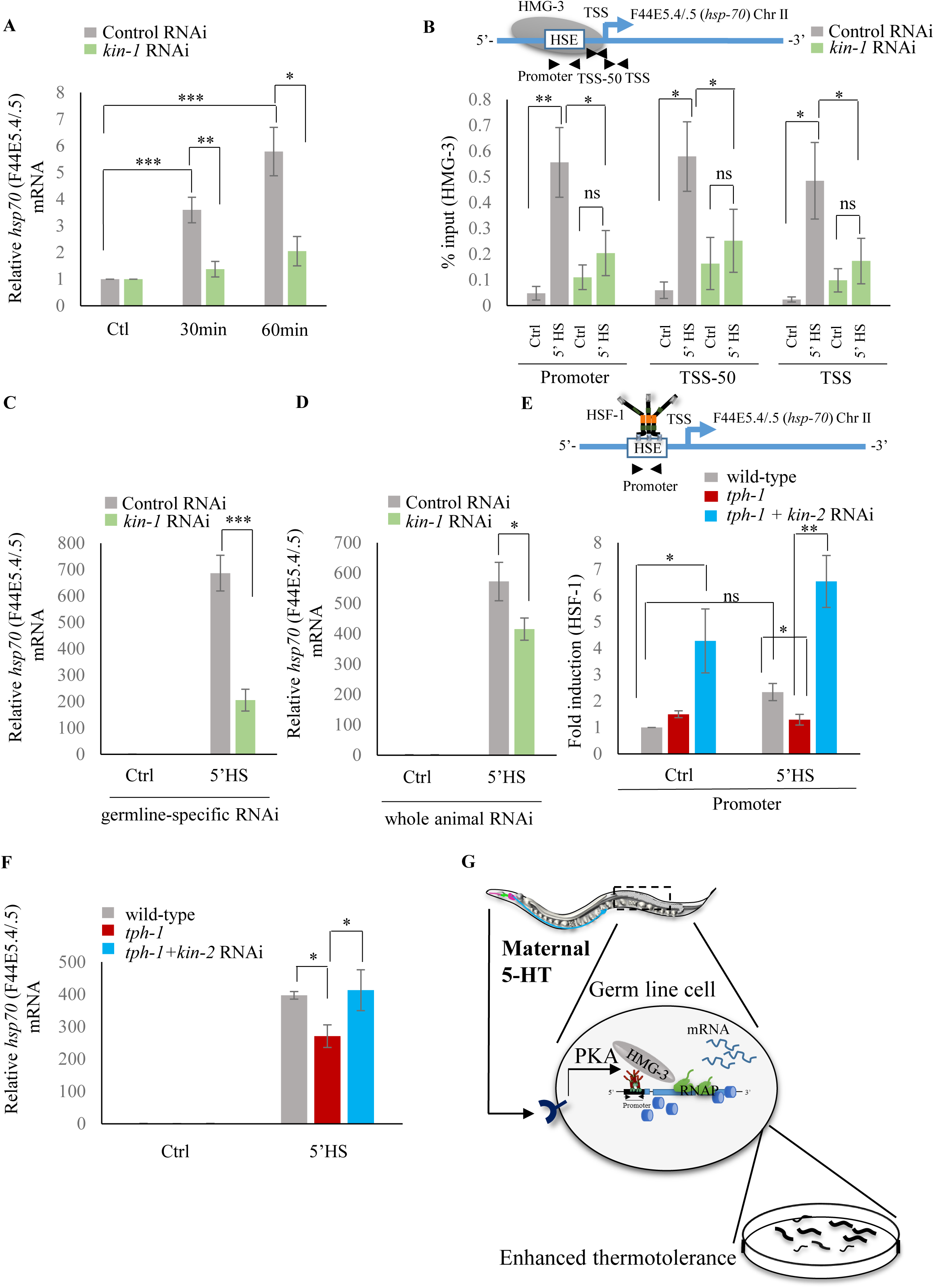
Serotonin-induced PKA-activation is a conserved signaling pathway that enables HSF-1 to recruit FACT in *C. elegans.* **A,** *hsp70* (F44E5.4/.5) mRNA levels in control-RNAi treated and *kin-1-*RNAi treated animals at different time points following optogenetic stimulation of 5-HT release. mRNA levels are normalized to control RNAi and *kin-1* RNAi treated unstimulated animals respectively (n=8 experiments). **B**, **Top:** Schematic of *hsp70* (F44E5.4/.5) gene showing Promoter, TSS-50 and TSS region used to assess HMG-3 occupancy. **Bottom:** HMG-3 occupancy (% input) in wild-type animals subjected to control and *kin-1* RNAi following 5 minutes at 34°C, across Promoter, TSS-50 and TSS regions (n=9 experiments). Specificity and efficiency of pull-down under control conditions was ascertained (see **Supplementary Figure 9**). **C**, *hsp70* (F44E5.4/.5) mRNA levels in control and heat-shocked (*mkcSi13* [*sun-1p::rde-1::sun-1 3′UTR* + *unc*-*119*(+)] *II; rde-1*(*mkc36*) *V*) animals that undergo germ-line specific RNAi following exposure to control RNAi or *kin-1* RNAi. HS: 5 minutes at 34°C (n=4 experiments). **D**, *hsp70* (F44E5.4/.5) mRNA levels in control and heat-shocked wild-type animals after they were subject to control and *kin-1* RNAi-mediated knockdown. HS: 5 minutes at 34°C (n=4 experiments). **E, Top**: Schematic of *hsp70* (F44E5.4/.5) promoter region assayed for HSF-1 occupancy. **Bottom:** HSF-1 occupancy in control and heat-shocked wild-type animals*, tph-1* mutant animals and *tph-1* mutant animals subjected to *kin-2* -RNAi. HS: 5 minutes at 34°C (n=5 experiments). % input values were normalized to that in control wild-type animals not subjected to heat shock. **F,** *hsp70* (F44E5.4/.5) mRNA levels in control and heat-shocked wild-type animals, *tph-1* mutant animals and *tph-1* mutant animals subjected to *kin-2* -RNAi. HS: 5 minutes at 34°C (n=5 experiments). Data show Mean ± Standard Error of the Mean. *, *p*<0.05; **, *p*< 0.01 ***, *p*<0.001; ns, non-significant. (**B, E**: ANOVA with Tukey’s correction, **A, C, D, F**: paired Student’s t-test). **G,** Working model showing how maternal 5-HT protects future progeny. 5-HT released by neurons permeates the animal through the coelomic fluid to act on the germline, activate the PKA signal transduction pathway in a tissue-autonomous manner and enable HSF-1 to recruit FACT, displace nucleosomes in the germline, and accelerate the onset of protective gene expression.

These data together allow us to propose a model whereby maternal 5-HT released by neurons acts on germ cell through 5-HT-mediated PKA signaling to hasten the timing of stress-gene expression by phosphorylating HSF-1, enabling it to recruit FACT, displace nucleosomes and promote RNAP transcription though chromatin. The activation of HSF-1 by PKA signaling occurs in the germline, as knocking down *kin-1* only in germ cells is enough to compromise *hsp* mRNA expression upon the 5 minutes heat-shock. Although we do not show it, we hypothesize that 5-HT likely acts directly on the germline cells upon release from neurons, due to its ability to diffuse through the coelomic fluid and bind 5-HT receptors throughout the animal. This, in turn leads to enhanced thermotolerance of the progeny of heat-shocked mothers (Figure 7G).

## Discussion

One of the more recent developments in the regulation of stress responses has been the demonstration that in *C. elegans* the activation of the unfolded protein response (UPR) in the cytoplasm^1,2,35^, endoplasmic reticulum^90^ and mitochondria^91^ are controlled cell non-autonomously by the nervous system. However, the mechanism by which this occurs was not known; neither was it clear whether such regulatory control was conserved. Here we show that 5-HT released from maternal neurons in *C. elegans* upon stress allows the information of the stress stimulus to be linked to the onset of protective HSF1-dependent transcription in germ cells, ensuring their survival upon fertilization and enhanced stress tolerance as larvae. 5-HT-mediates these effects by enabling HSF-1 to modify chromatin through the activity of the histone chaperone FACT and accelerate the onset of transcription by displacing histones. Thus, 5-HT release by neurons, in effect, sets the level of the physiological stimulus required to activate a transcriptional response amongst the germline nuclei. Remarkably, maternal 5-HT release upon stress causes an increase in *hsp70* mRNA levels in embryos and transgenerational stress tolerance in progeny. Given the role of neuronal 5-HT in modulating memory and learning these studies have wide-ranging implications for the effects on maternal experience on progeny physiology. Although the exact mechanism by which embryos have more *hsp* mRNA was not explored, this is consistent with published observations that even if oocytes are not transcriptionally competent, pachytene nuclei can act as nurse cells to provide material to uncellularized oocytes^92^ to be subsequently utilized by embryos. It is possible that besides *hsps,* 5-HT also stimulates the packaging of other mRNAs into soon-to be fertilized embryos to promote their survival. This remains to be determined. In mammalian cells 5-HT activates HSF1 through the 5-HT4 receptor. *C. elegans* do not possess a direct ortholog of 5-HT4^23,24^ and the receptor required to activate PKA in the germline remains to be identified. One possible receptor is the SER-7, which although has not been localized to the germline, controls numerous aspects of egg-laying and acts through a Gα_s_-coupled signaling pathways to promote PKA-dependent phosphorylation of target proteins^23,24^.

Our data also show that for unknown reasons, fertilized *C. elegans* embryos are exquisitely vulnerable to even transient temperature fluctuations. Lowering of the threshold of transcription onset in the germline by 5-HT is therefore critical to protect development, and indeed the germline is amongst the first tissues to express protective *hsp70s*. Across tissues and during development, transcriptional responses display characteristic dynamics and thresholds of activation that are linked to their biological function^31,61,93,94^. We provide here a molecular mechanism by which thresholds for activation of transcription upon stress can be set to different levels in different tissues *in vivo.* The essential aspects of the 5-HT signaling pathway are conserved in mammalian neurons. Moreover, the involvement of PKA in 5-HT dependent HSF1 activation is intriguing given the role of both 5-HT and PKA in cellular plasticity. It is therefore tempting to speculate that the ability to modulate transcriptional dynamics through modifying chromatin accessibility may be a more general function of 5-HT in neurodevelopment and as a neuromodulator, allowing it to steer developmental timing and neuronal activity^16, 19–21,95^. In addition, this ability to functionally activate HSF1 in mammalian neurons and human cells, in the absence of proteotoxicity through activation of 5HT4 receptors, could have implications for the treatment of neurodegenerative diseases where HSF1 is protective^33,96,97^.

Why might 5-HT, an abundant neuromodulator and signaling molecule that portends growth, also modulate stress responsiveness of germ cells? The answer to this question may be limited by our understanding of what precisely constitutes ‘stress’ to different cells. Germ cells typically consist of ‘poised’ chromatin^98^ bearing both activation and repressive histone marks, which potentially can resolve into growth-related and ‘active’, or stress-related and ‘repressed’ antagonistic gene expression programs^99^. Across the animal kingdom, 5-HT release can signal stress or growth ^6,8^. We postulate that the ability of 5-HT to modulate chromatin accessibility in response to environmental input may allow it to function as a switch at the nexus of these essential programs.

## Supplementary Figure Legends

**Supplementary Figure 1:**
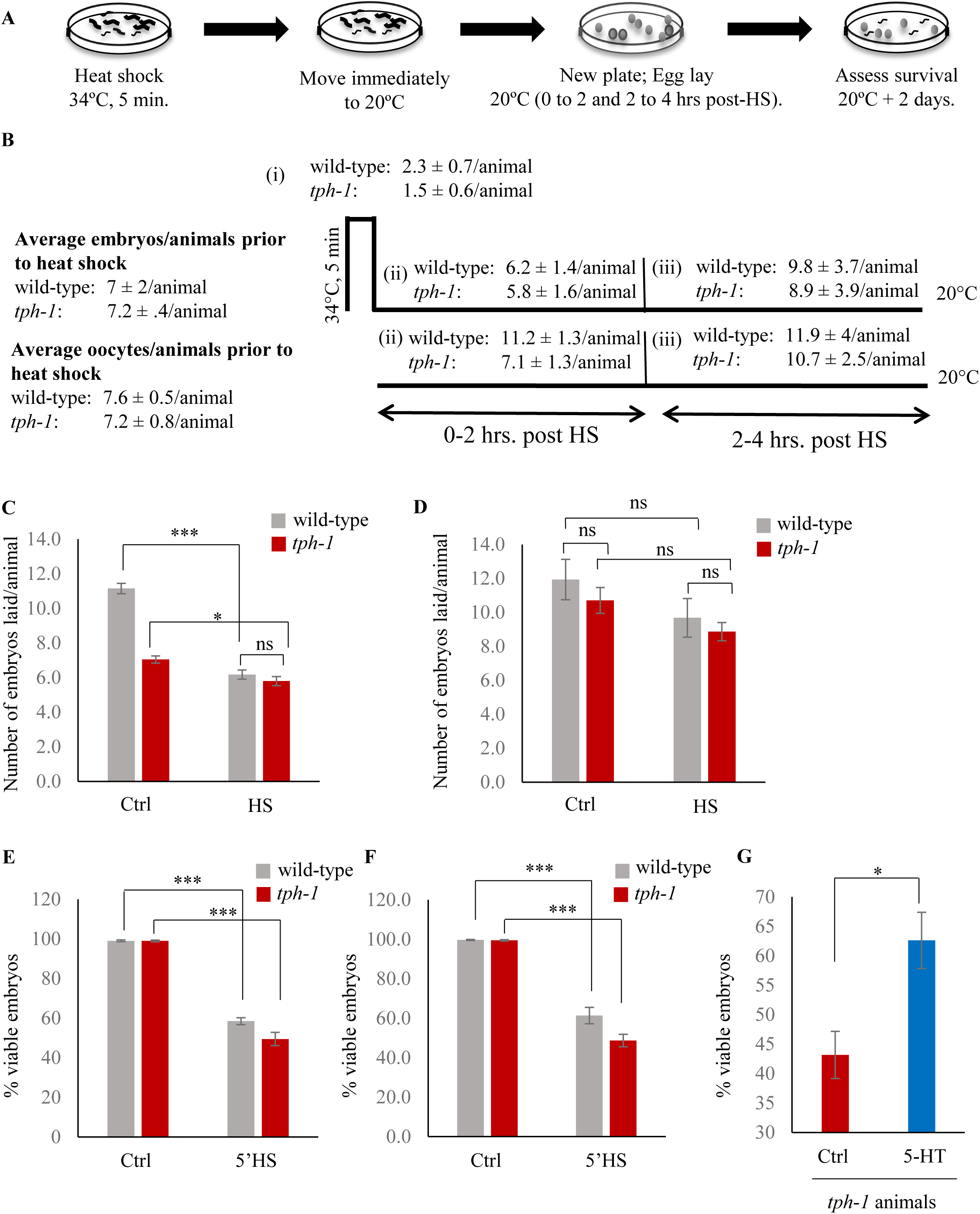
Characterizing the effects of heat on viability of embryos. **A,** Schematic design of the experiment. **B,** The average numbers of embryos and oocytes present per animal in wild-type animals and *tph-1* mutants prior to heat exposure, and embryos laid per animal under control conditions at 20°C, (i), upon the 5 minute exposure to 34°C, (ii), during 0-2 hours post heat shock, and (iii) during 2-4 hours post heat shock are shown. **C,** Mean numbers of embryos laid per animal by wild-type animals and *tph-1* mutants under control conditions at 20°C, and during 0-2 hours post-heat shock (n=18 experiments). **D,** Mean numbers of embryos laid per animal by wild-type animals and *tph-1* mutants under control conditions, and 2-4 hours post heat shock. (n=28 experiments). **E,** Percent viable embryos when embryos laid by wild-type animals and *tph-1* mutant animals during a 2 hour interval were exposed to 34°C for 5 minutes. **F,** Percent viable embryos when embryos were present *in utero* as wild-type animals or *tph*-*1* mutant animals were exposed to 34°C for 5 minutes. This was determined by assaying the viability of embryos laid during a 2 hour interval immediately after the mothers were heat-shocked. **G,** Percent viable embryos laid by *tph-1* mutant animals upon treatment with exogenous 5-HT compared to those laid by untreated *tph-1* mutant animals. Data in **C-G** show Mean ± Standard Error of the Mean. *, *p*<0.05; **, *p* < 0.01 ***, *p*<0.001, ns=non-significant; (Paired Student’s t-test). ns, non-significant.

**Supplementary Figure 2:**
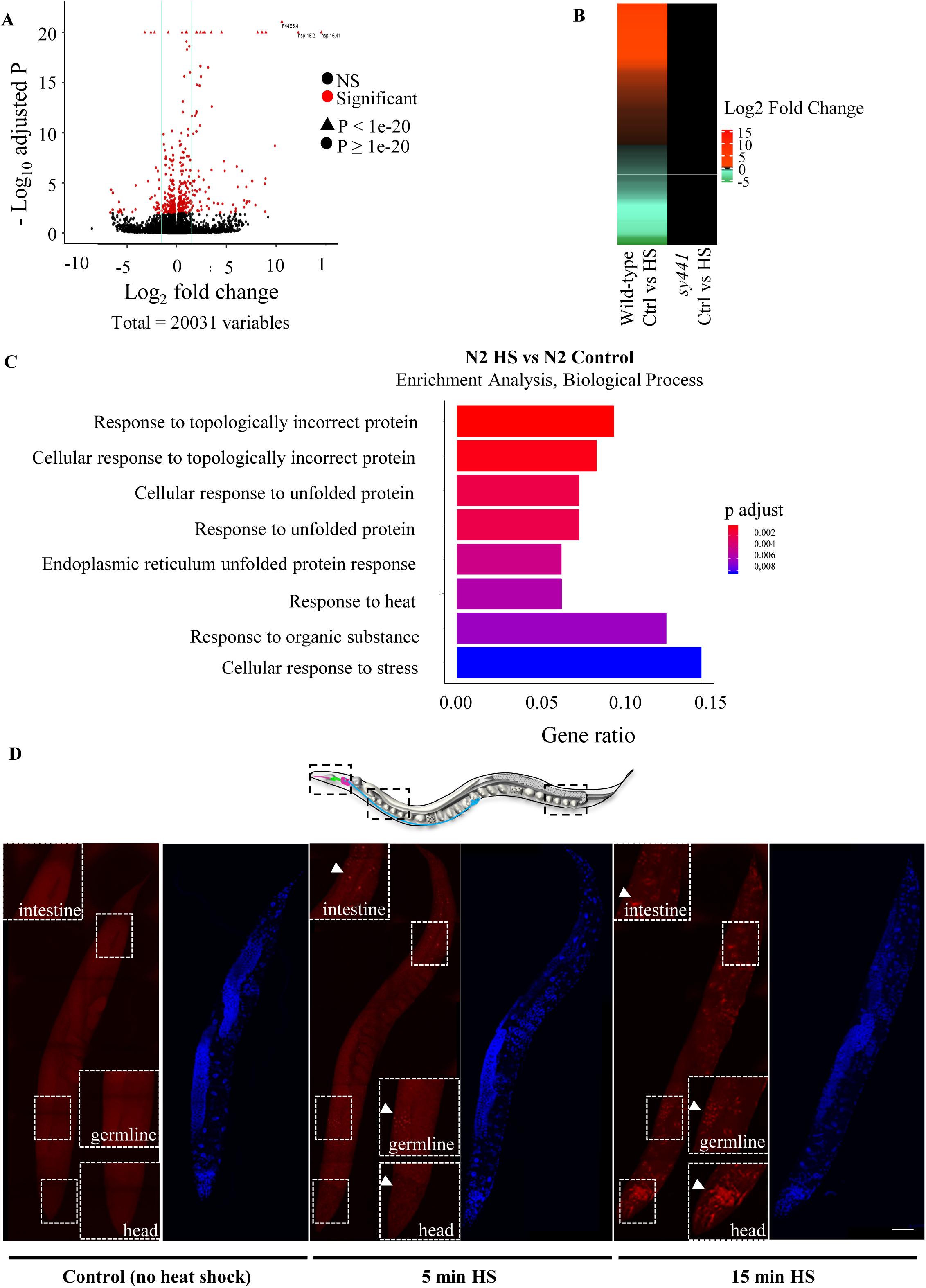
A brief (5-minute) heat-shock induces HSF-1-dependent gene expression. **A,** Volcano plot from RNA-seq data showing differentially expressed transcripts (Log2 fold change) in wild-type animals following 5 minutes heat shock at 34°C compared to control, wild-type animals. Genes differentially expressed at 0.01 FDR are in red. **B,** Heat map showing differential gene expression in *hsf*-*1(sy441)*I mutants (Log2 fold change) that lack functional HSF-1 upon heat shock, compared to differential gene expression in wild-type animals exposed to the same conditions . Heat shock: 5 minutes heat shock at 34°C. **C**, Gene Ontology analysis (Biological Processes) of differentially expressed transcripts in wild-type animals following a 5 minute heat shock at 34°C. **D,** Representative micrographs showing projections of confocal images of wild-type animals processed for *hsp70* (F44E5.4/.5) mRNA localization using smFISH. *hsp70* (F44E5.4/.5) mRNA localization under control conditions and upon 5-minute and 15-minute exposure to 34°C (n=5 experiments, 33 animals). Red.: *hsp70* (F44E5.4/.5) mRNA. Blue: DAPI stained nuclei. Arrows indicate *hsp* mRNA. Insets correspond to specific regions within the animal. Scale bar=10µm

**Supplementary Figure 3:**
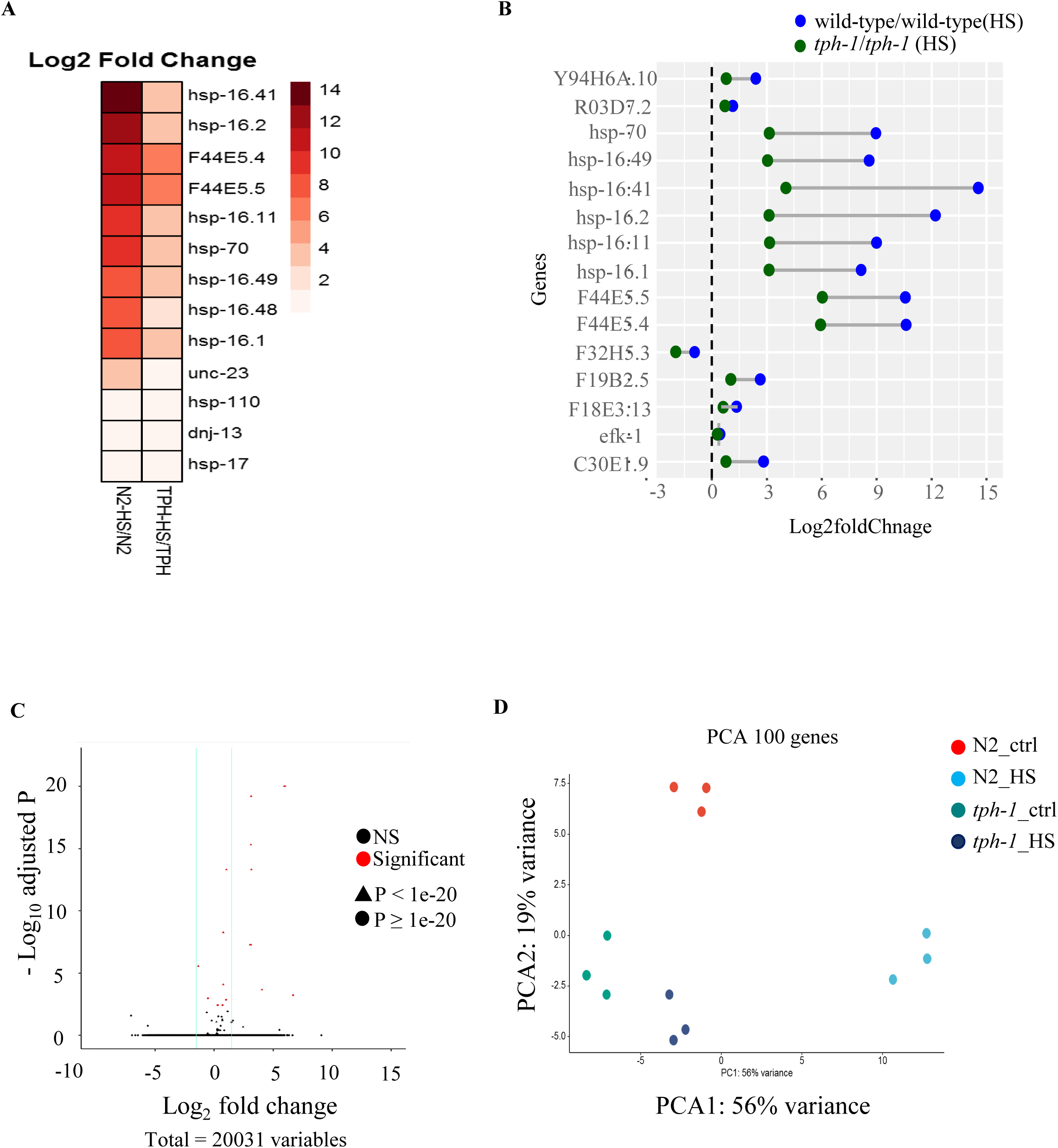
The lack of serotonin diminishes HSF-1-dependent gene expression upon 5-minute heat-shock. **A,** Heat map depicting expression levels (Log2 fold change) of thirteen differentially expressed *hsp* genes in wild-type and *tph-1* mutants. **B,** Dumb-bell graph showing the expression levels of the 15 differentially expressed genes that are common between wild-type and *tph-1* heat shock response. Heat shock: 5 minutes heat shock at 34°C. **C,** Volcano plot from RNA-seq data showing differentially expressed transcripts (Log2 fold change) in *tph-1* mutant animals following 5 minutes heat shock at 34°C compared to control, *tph-1* animals. Compare with Supplementary Figure 2A. **D,** Principal Component Analysis (PCA) depicting the separation of wild-type animals exposed for 5 minutes to 34°C from control non-heat shocked animals, and the lack of similar separation in *tph-1* mutant animals.

**Supplementary Figure 4:**
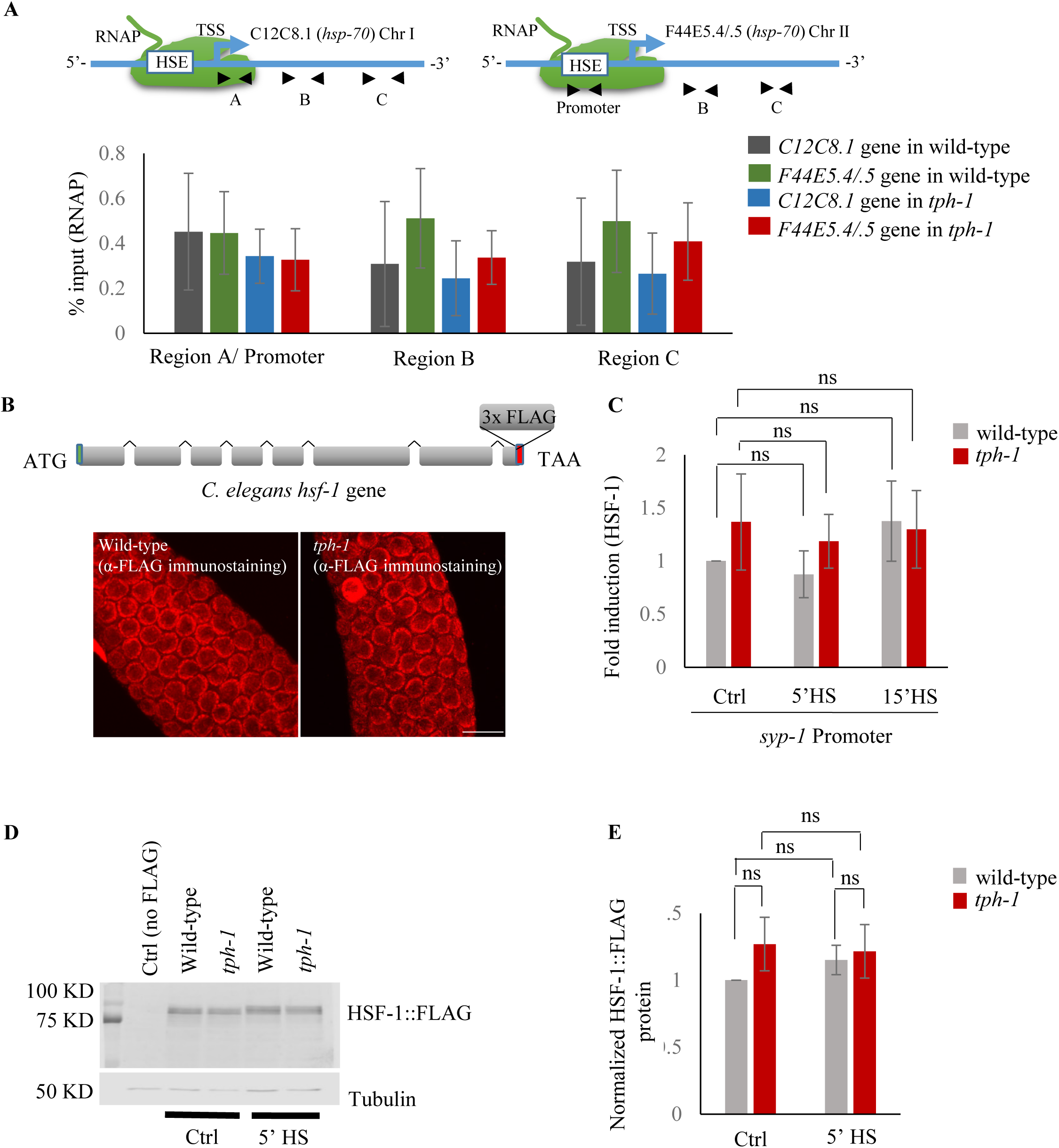
Characterization of RNAP and HSF-1 for ChIP-qPCR assays. **A, Top:** ChIP-qPCR to measure RNAP occupancy: Schematic of *hsp70* (F44E5.4/.5) gene regions within the Promoter (−390 to −241), middle of gene (Region B: +696 to +915) and towards 3’-UTR (Region C: +1827 to +1996) assayed for occupancy by RNAP. Schematic of *hsp70* (C12C8.1) gene regions close to the beginning (Region A: +25 to +185), middle of gene (Region B: +475 to +583) and towards 3’-UTR (Region C:+1645 to +1835) assayed for occupancy by RNAP. **Bottom:** RNAP occupancy at these region in control non-heat shocked wild-type and *tph-1* mutant animals. RNAP (% input) at the RegionA/Promoter region in control wild-type animals does not significantly differ from that at Region B or Region C, as would have been expected if RNAP was paused near the TSS (n=8 experiments). Data show Mean ± Standard Error of the Mean. ANOVA with Tukey’s correction**. B, Top:** Schematic of *hsf-1* tagged at the endogenous locus with 3X FLAG using CRISPR/Cas9. **Bottom:** Representative micrographs of confocal sections of the germline region showing HSF-1 immunostaining with anti-FLAG antibody in wild-type and *tph-1* mutants (n=5 experiments). Note HSF-1 is expressed in germline cells in both wild-type animals and *tph-1* mutant animals. Scale bar=10µm. **C,** Fold change in HSF-1 occupancy at the promoter of a control gene not having any HSF-1 binding sites (*syp-1*) in wild-type animals and *tph-1* mutants following 5 and 15 minutes at 34°C. % input values were normalized to that in control wild-type animals (n=14 experiments). **D,** Representative Western blot using anti-FLAG antibody showing the specificity of the FLAG antibody (lane 1: Ctrl are wild-type animals that do not express FLAG-tagged proteins), and HSF-1 levels in wild-type and *tph-1* animals under control and heat shock conditions. Tubulin was used as a loading control. **E,** Quantified HSF-1 levels in wild-type and *tph-1* animals under control and heat shock conditions (n=3 experiments). Protein levels are normalized to control wild-type animals. Note HSF-1 protein levels do not change. Data in **A, C, E** show Mean ± Standard Error of the Mean. **A** and **C**, ANOVA with Tukey’s correction. **E,** paired Student’s t-test. ns, non-significant.

**Supplementary Figure 5:**
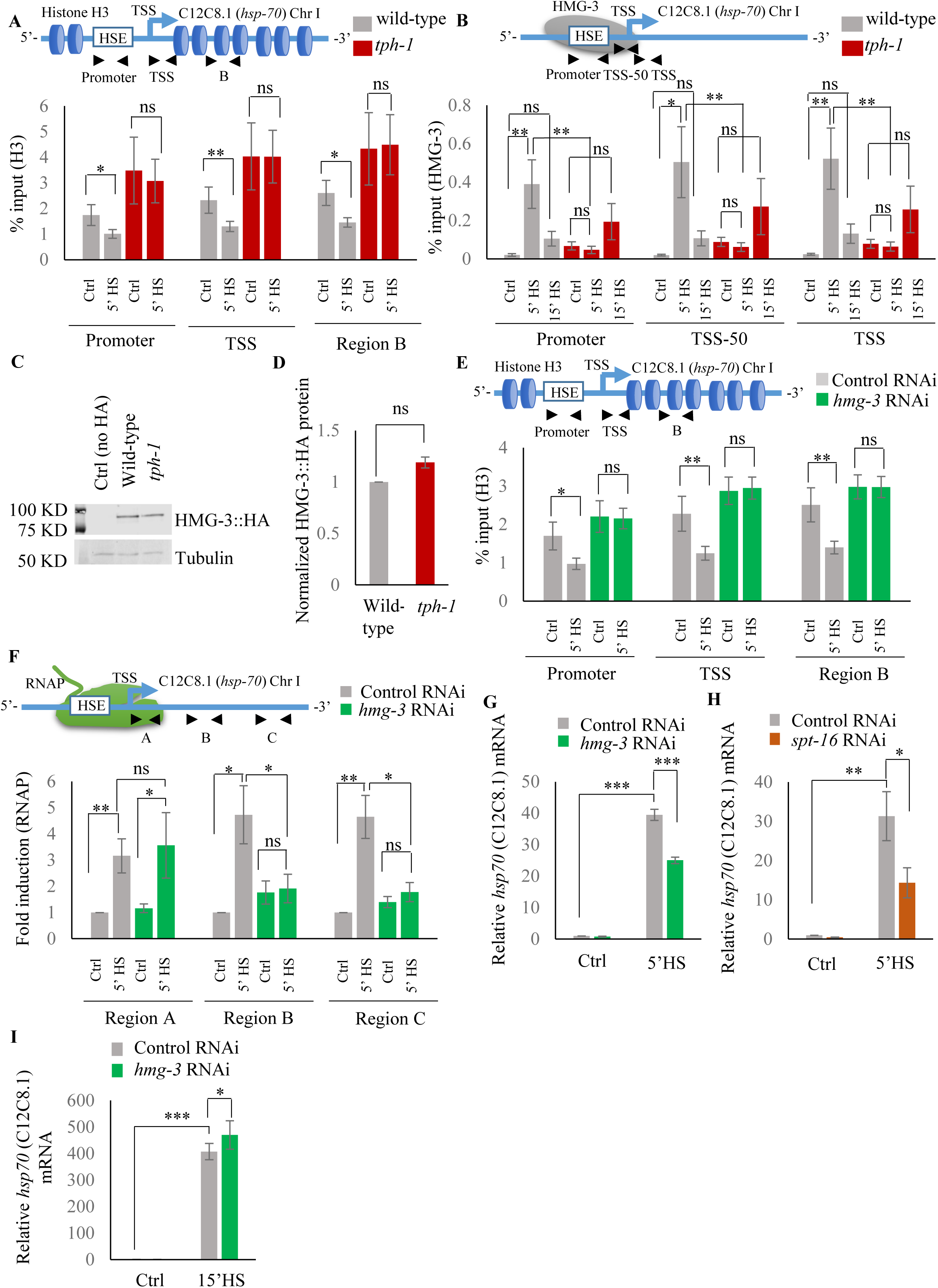
Characterizing H3 and FACT occupancy across *hsp* genes in wild-type and *tph-1* mutant animals under control and heat shock conditions. **A, Top:** Schematic of *hsp70* (C12C8.1) gene regions within the Promoter (same as in Figure 3D: -166 to --78), Transcription Start Site (TSS: +40 to +131) and gene body (Region B: +475 to +583) assayed for histone H3 occupancy. **Bottom:** Occupancy of histone H3 (% input) at the Promoter, TSS and Region B upon 5 minutes at 34°C (n=9 experiments). **B, Top:** Schematic of *hsp70* (C12C8.1) gene regions within the Promoter (same as Figure 3D: -166 to –78), region upstream of Transcription Start Site (TSS-50: -77 to +42) and Transcription Start Site (TSS: +40 to +131) assayed for HMG-3 occupancy in wild-type and *tph-1* mutant animals. **Bottom:** HMG-3 occupancy (% input) across Promoter, TSS-50 and TSS following 5 and 15 minutes at 34°C (n=13 experiments). The specificity and efficiency of pull-down under control conditions was ascertained. **C,** Representative Western blot using anti-HA antibody showing the specificity of the HA antibody (lane 1: Ctrl are wild-type animals that do not express HA-tagged proteins), and HMG-3 protein levels in wild-type and *tph-1* animals under control conditions. Tubulin was used as the internal control. **D,** Quantified levels of HMG-3 in wild-type and *tph-1* animals (n=3 experiments). Protein levels were normalized to that in control animals. **E, Top:** Schematic of *hsp70* (C12C8.1) gene: same regions as in **A** were assayed for H3 occupancy. **Bottom:** Occupancy of histone H3 (% input) in Promoter, TSS and in Region B of *hsp70* (C12C8.1) under control conditions and following 5 minutes at 34°C, in control-RNAi treated animals and *hmg-3-*RNAi treated animals (n=9 experiments). **F, Top:** Schematic of *hsp70* (C12C8.1) gene: regions as in Figure 3C, i.e. close to the beginning (Region A: +25 to +185), middle of gene (Region B: +475 to +583) and towards 3’-UTR (Region C:+1645 to +1835) were assayed for occupancy by RNAP following *hmg-3* knock-down by RNAi. **Bottom:** Fold change in RNAP at beginning, Region B and Region C following 5-minute heat shock at 34°C (n=5 experiments). % input values were normalized to that in control-RNAi treated animals at these regions. **G,** *hsp70* (C12C8.1) mRNA levels in control-RNAi treated and *hmg-3* -RNAi treated animals following a 5-minute heat shock at 34°C (n=6 experiments). **H,** *hsp70* (C12C8.1) mRNA levels in control-RNAi treated and *spt-16* -RNAi treated animals following a 5-minute heat shock at 34°C (n=4 experiments). **I,** *hsp70* (C12C8.1) mRNA levels in control-RNAi treated and *hmg-3* -RNAi treated animals following a 15-minute heat shock at 34°C (n=4 experiments). Data show Mean ± Standard Error of the Mean. *, *p*<0.05; **, *p*< 0.01 ***, *p*<0.001; (paired Students t-test). **A, B, E, F**: ANOVA with Tukey’s correction. **D**, **G-I**: paired Student’s t-test. ns, non-significant.

**Supplementary Figure 6:**
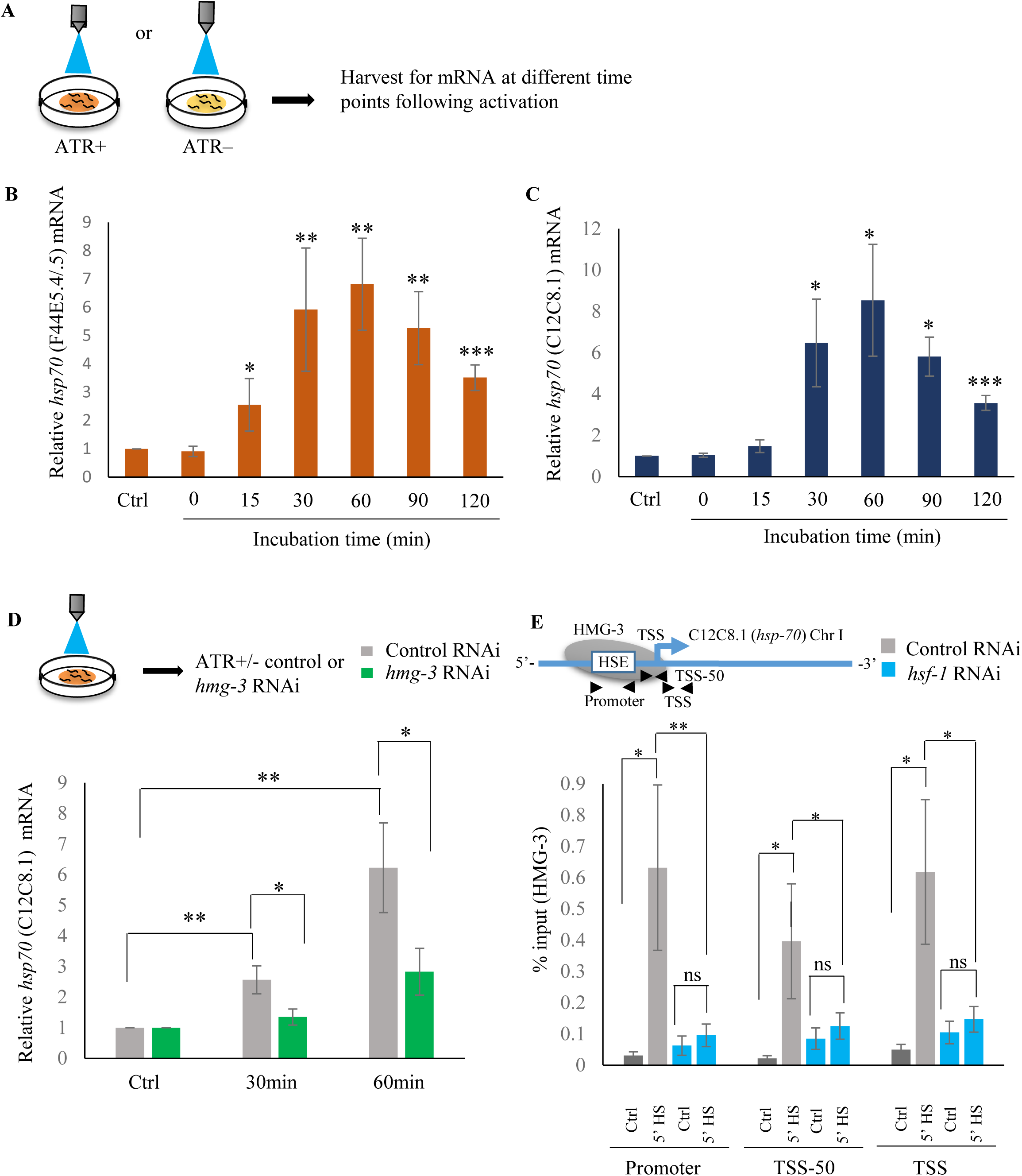
Serotonin-induced transcriptional activity of HSF-1 in *C. elegans* is FACT dependent. **A,** Schematic of optogenetic activation of 5-HT release by stimulating ADF and NSM neurons. **B, C,** *hsp70* (F44E5.4/.5) and *hsp70* (C12C8.1) mRNA levels respectively, at different time points following optogenetic stimulation (n=6 experiments)**. D, Top:** Schematic of optogenetic activation of 5-HT release by stimulating ADF and NSM neurons. **Bottom:** *hsp70* (C12C8.1) mRNA levels in control-RNAi treated animals and animals treated with *hmg-3-*RNAi following optogenetic activation of 5-HT release. mRNA levels were normalized to control-RNAi treated and *hmg-3*-RNAi treated unstimulated animals (n=6 experiments). **E Top:** Schematic of *hsp70* (C12C8.1) gene showing Promoter, TSS-50 and TSS to assess HMG-3 occupancy in control-RNAi treated and *hsf-1-*RNAi treated animals. **Bottom:** HMG-3 occupancy (% input) across Promoter, TSS-50 and TSS in control-RNAi and *hsf-1* -RNAi treated animals following 5 minutes at 34°C (n=9 experiments). Specificity and efficiency of pull-down under control conditions was ascertained. Data show Mean ± Standard Error of the Mean. *, *p*<0.05; **, *p*< 0.01 ***, *p*<0.001; (**B-D**, Paired Student’s t-test. **E**, ANOVA with Tukey’s correction). ns, non-significant.

**Supplementary Figure 7:**
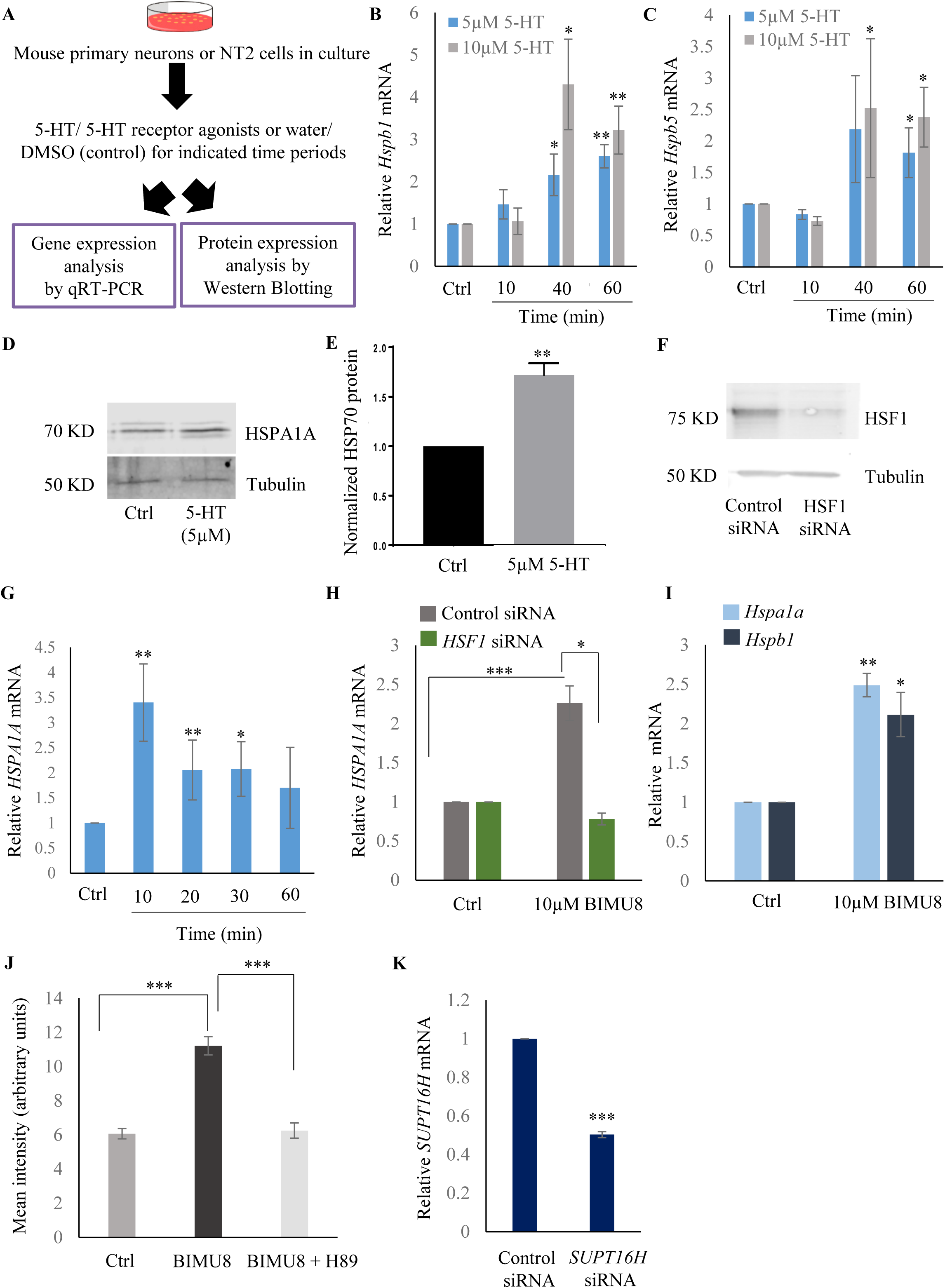
Serotonin activates a PKA-mediated signal transduction pathway to enable HSF1-FACT interaction in mammalian cells. **A,** Experimental setup for assessing whether 5-HT is a conserved signal that activates HSF1 in mammalian cells. **B, C,** Time and dose-dependent change in *Hspb1* and *Hspb5* mRNA levels respectively, in control and 5-HT treated primary cortical neuronal cultures (n=4 experiments). **D,** Representative Western blot showing HSPA1A protein levels following 5-HT stimulation of NT2 cells (5µM was applied for 24 hrs to be able to assess protein accumulation; n=4 experiments). **E,** Quantitation of HSPA1A protein levels in control and 5-HT treated NT2 cells (n=3 experiments). Tubulin was used as internal control. **F,** Representative Western blot showing HSF1 protein levels in control and *HSF1* siRNA treated NT2 cells to confirm siRNA knockdown of HSF1 (n=2 experiments). **G,** Temporal dynamics of *HSPA1A* mRNA levels in control and BIMU8 treated NT2 cells (n=5 experiments). **H,** *HSPA1A* mRNA levels in untreated and BIMU8 treated NT2 cells (10 µM for 10 minutes) transfected with control and *HSF-1* siRNA (n=4 experiments). **I,** *Hspa1* and *Hspb1* mRNA levels in untreated and BIMU8 treated (10 µM for 10 minutes) primary cortical neuronal cultures (n=3 experiments). **J,** Quantitation of fluorescence intensity following immunostaining for HSF1 in the nuclei of untreated NT2 cells, NT2 cells treated with BIMU8 (10 µM for 10 minutes), and NT2 cells treated with BIMU8 and H89. Fluorescence intensity values (arbitrary units) were derived using ImageJ following background subtraction. (n=2 experiments; 25 cells). **K,** *SUPT16H* mRNA levels in control-siRNA and *SUPT16H-*siRNA treated NT2 cells to confirm siRNA knockdown of *SUPT16H* (n=5 experiments). Data in **B, C, E, G-K** show Mean ± Standard Error of the Mean. Values were normalized to that in either control untreated cells, or control-siRNA treated cells. *, *p*<0.05; **, *p*< 0.01 ***, *p*<0.001; (paired Students t-test).

**Supplementary Figure 8:**
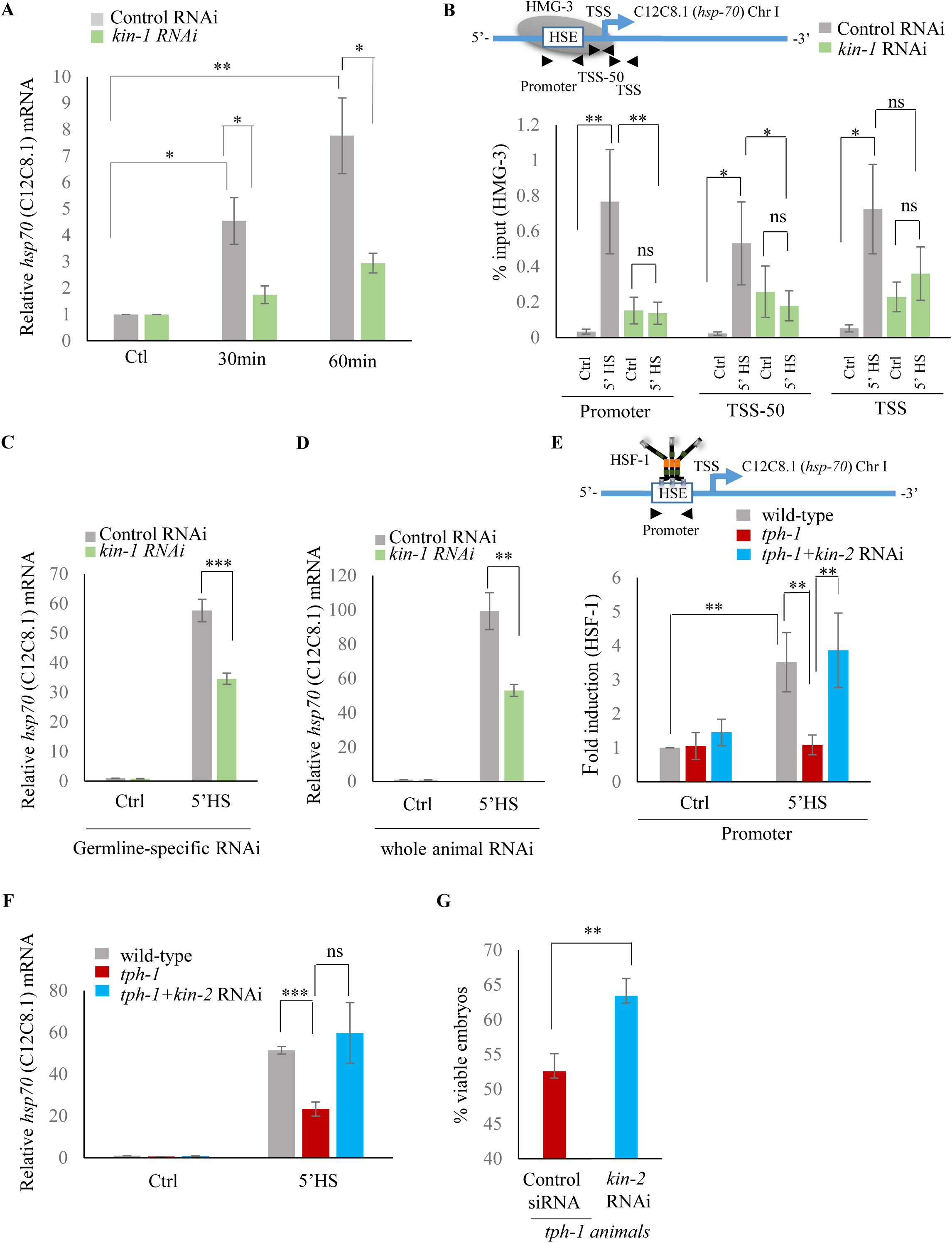
Serotonin-induced PKA-activation is a conserved pathway that enables HSF-1 to recruit FACT in *C. elegans*. **A,** *hsp70* (C12C8.1) mRNA levels in control RNAi treated and *kin-1* RNAi treated animals collected at different time points following optogenetic stimulation to release 5-HT. mRNA levels were normalized to control RNAi treated or *kin-1* RNAi treated unstimulated animals (n=8 experiments). **B, Top:** Schematic of *hsp70* (C12C8.1) gene showing Promoter, TSS-50 and TSS regions used to assess HMG-3 occupancy. **Bottom:** HMG-3 occupancy (% input) in wild-type animals subjected to control-RNAi and *kin-1-*RNAi following a 5-minute heat shock treatment at 34°C (n=9 experiments). **C,** *hsp70* (C12C8.1) mRNA levels in control and heat-shocked (*mkcSi13* [*sun-1p::rde-1::sun-1 3′UTR* + *unc-119*(+)] *II; rde-1*(*mkc36*) *V*) animals that undergo germline specific RNAi; animals were subject to control and *kin-1* RNAi-mediated knockdown. HS: 5 minutes at 34°C (n=4 experiments). **D**, *hsp70* (C12C8.1) mRNA levels in control and heat-shocked animals wild-type animals after they were subject to control and *kin-1* RNAi-mediated knockdown. HS: 5 minutes at 34°C (n=4 experiments). **E, Top**: Schematic of *hsp70* (C12C8.1) promoter region assayed for HSF-1 occupancy. **Bottom:** HSF-1 occupancy at the *hsp70* (C12C8.1) gene in control and heat-shocked wild-type animals, control and heat-shocked *tph-1* mutant animals, and control and heat-shocked *tph-1* mutant animals subjected to *kin-2* -RNAi. HS: 5 minutes at 34°C (n=5 experiments). % input values were normalized to that in control wild-type animals not subjected to heat shock. **F,** *hsp70* (C12C8.1) mRNA levels in control and heat-shocked animals wild-type animals, *tph-1* mutant animals and *tph-1* mutant animals subjected to *kin-2* -RNAi. HS: 5 minutes at 34°C (n=5 experiments). **G,** Percent viable embryos laid 2-4 hrs. post-heat shock by *tph-1* animals and *tph-1* animals subjected to *kin-2* RNAi (n=5 experiments). Data show Mean ± Standard Error of the Mean *, *p*<0.05; **, *p*< 0.01 ***, *p*<0.001; (**B, E**: ANOVA with Tukey’s correction, **A, C, D, F**, G: paired Student’s t-test). ns, non-significant.

**Supplementary Figure 9:**
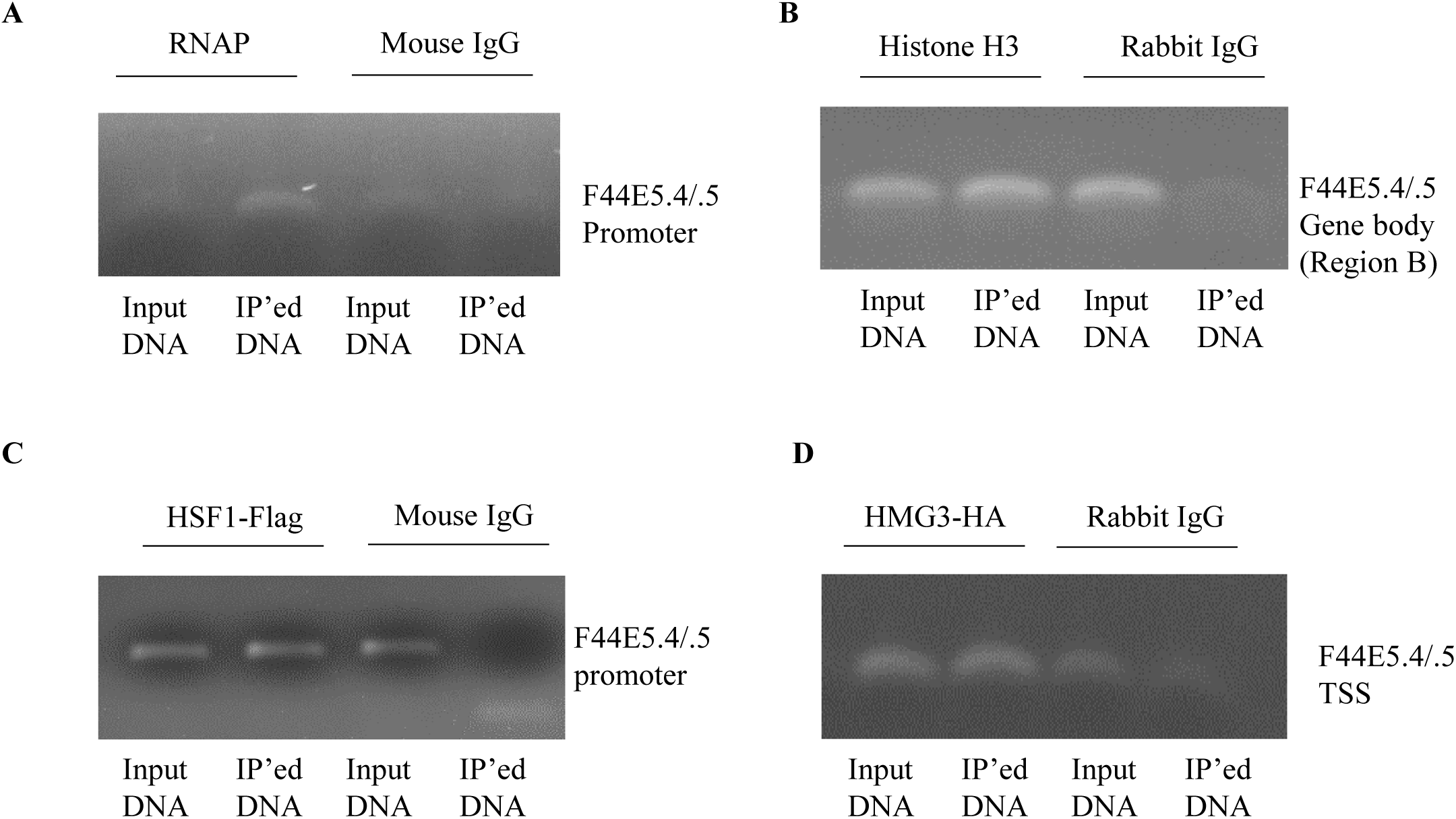
Validation of specificity and efficiency of antibodies used for ChIP assays. **A-D,** Representative agarose gels used to assess specificity and efficiency of immunoprecipitation (IP) and primer sequences used for ChIP-PCR. All amplified bands were the expected size, mouse and rabbit control IgG did not yield detectable signals. Most importantly, we confirmed that the ChIP signal in control animals was detectable and real, to validate the calculation of normalized fold changes in our experiments. **A**: IP from wild-type control animals using mouse antibody to total RNAP. Mouse IgG used as control. Primers at the Promoter region of *hsp70* (F44E5.4/.5). **B**: IP from wild-type control animals using rabbit antibody to histone H3. Rabbit IgG used as control. Primers in the Region B of *hsp70* (F44E5.4/.5). **C**: IP from wild-type control animals using mouse anti-FLAG antibody. Mouse IgG used as control. Primers at the Promoter region of *hsp70* (F44E5.4/.5). **D**: IP from wild-type control animals using rabbit anti-HA antibody. Rabbit IgG used as control. Primers at the TSS region of *hsp70* (F44E5.4/.5).

## Acknowledgements

We thank the members of V.P. laboratory, Dr. Sarit Smolikove, Dr. Bin He, Dr. Chris Stipp and Dr. Tali Gidalevitz for their helpful comments, Kat Dvorak, Matthew Wheat, Dr. Rachel Reichman and Gery Hehman for technical help, and Dr. Smolikove for advice with CRISPR/Cas9. The BAT1560 strain was a kind gift from Dr. Baris Tursun, Max Delbrück Center (MDC). We would also like to thank anonymous reviewers of a prior submission for extremely helpful comments. HSP70 antibodies were a kind gift of Dr. Morimoto (Northwestern University). Nematode strains were provided by the Caenorhabditis Genetics Center (CGC) (funded by the NIH Infrastructure Programs P40 OD010440). We acknowledge the CCG and the Aging Mind and Brian Initiative (AMBI) at University of Iowa for support. This work was supported by NIH R01 AG 050653 (V.P.).

## Methods

### C. elegans strains

Most *C. elegans* strains were obtained from Caenorhabditis Genetics Center (CGC, Twin Cities, MN). The BAT1560 strain was a kind gift from Dr. Baris Tursun, Max Delbrück Center (MDC) ^1^. The HSF-1::FLAG strain was created using CRISPR/Cas9.

| Strain Name | Genotype | Source |
| --- | --- | --- |
| HSF-1::FLAG | *hsf-1::flag I* | Prahlad lab |
| Wild-type | Bristol N2 | CGC |
| MT15434 | *tph-1 (mg280) II* | CGC |
| PS3551 | *hsf-1(sy441)I* | CGC |
| AQ2050 | *lite-1(ce314);_ljIs102 [tph-1;;ChR2::YFP;unc-122::gfp]* | CGC |
| BAT1560 | *hmg-3(bar24[hmg-3::3xHA]) I* | Dr. Baris Tursun, Max Delbrück Center (MDC) |
| SS104 | *glp-4(bn2) I* | CGC |
| NL2098 | *rrf-1(pk1417) I* | CGC |
| DCL569 | *mkcSi13 II; rde-1(mkc36) V* | CGC |
| VP303 | *rde-1(ne219) V; kbIs7* | CGC |

#### Generation of *hsf-1*::FLAG

CRISPR/Cas9 was used to create *C. elegans* strains where the endogenous *hsf-1*(I) gene was tagged at the C terminus with a 3X FLAG sequence to create HSF-1::FLAG animals. Individual adult worms were injected on 3% agarose pads with the injection mix detailed below. Following injection, animals were singled onto NGM plate. Plates were screened for the rol or dpy phenotypes created by the co-CRISPR marker *dpy-10*^2^ . One hundred animals with a DPY or Roller phenotype were isolated as F1s and screened for the FLAG insertion by PCR. Three days later, single wild-type F2 offspring from plates with Dpy and/or Rol offspring were singled, screened for homozygosity of the FLAG insertion by PCR, and sequenced. The Cas9 enzyme, ultramer oligonucleotides, tracrRNA, and crRNAs were obtained from IDT

|  |  |
| --- | --- |
| tracr RNA | 30 μM |
| crRNA- dpy10 | 5 μM |
| crRNA- TARGET | 25 μM |
| IDT Cas9 Enzyme | 12.2 μM |
| dpy-10 ssODN | 0.5 μM |
| Target ssODN | 5 μM |

The sequences of the crRNA and the ssODN for hsf-1 are:

**crRNA-C’:** AAGTCCATCGGATCCTAATT

**ssODNhsf-1-C:**

TCCCACATTCACCGGCTCTTCGTACTCCAAGTCCATCGGATCCTAATTTGGTTgactacaaagaccatgacggtgattataaagatcatgaTatcgaTtacaaggatgacgatgacaagTAAttattgattttttttttgaacgtttagctcaaaattcctctc

#### Generation of *tph-1*; *hsf-1*::FLAG and *tph-1*; *hmg-3*::HA

The *tph-1 (mg280) II* strain was crossed into the *hsf-1::FLAG*(I) strain or the *hmg-3::*HA (I) and verified by PCR.

### Growth conditions of *C. elegans* strains

All strains except *glp-4(bn2) I* were grown and maintained at 20°C, except *glp-4 (bn2) I* worms were grown and maintained at 15°C (permissive temperature). For the experiments involving *glp-4 (bn2) I*, *glp-4 (bn2) I* eggs or wild-type eggs were raised at the permissive temperature (15°C), or shifted to 25°C after they were laid at 15°C until animals were day-1 adults. Animals were grown and maintained at low densities in incubators under standard conditions by passaging 8-10 L4s onto nematode growth media (NGM) plates and, 4 days later, picking L4 animals onto fresh plates for experiments. Animals were fed *Escherichia coli* OP50 obtained from CGC that were seeded onto culture plates 2 days before use. The NGM plates were standardized by pouring 8.9 ml of liquid NGM per 60 mm plate weighed before use. Plates had an average weight of 13.5±0.2 g. Any plates that varied from these measurements were discarded. Ambient temperature was maintained at 20°C to 22°C and carefully monitored throughout the experimental procedures. All animals included in the experiments, unless otherwise stated, were 1-day-old hermaphrodites that were age-matched either by (a) bleaching and starting the experiment after 75-78 hours or (b) picking as L4 juveniles 24 to 26 hours before the start of the experiment.

### Mammalian cell culture

Dr. Christopher Stipp, University of Iowa, gifted NTERA-2 cl.D1 (also known as NT2) cells. For regular maintenance, cells were cultured in DMEM (Life Technologies) supplemented with 10% fetal bovine serum (Life Technologies), 2mM L-glutamine and 100U/ml penicillin and 100 µg/ml streptomycin. Cells were maintained at 37°C in 5% CO_2_ atmosphere under humidified conditions. Cell were passaged by splitting them (1:4) when cell confluence reached ∼90%. All cells used were between passage numbers 15-20. Cells were routinely checked for mycoplasma contamination.

### Mouse strain and cortical neuron culture

Cortical neuron cultures were performed essentially as described previously ^3,4^using P0 pups from timed-pregnant C57BL/6 mice (Harlan). Briefly, cortices were dissected, meninges were removed, and ∼1 mm^2^ pieces were digested in an enzyme solution (papain, 10 units/ml) 2 × 20 min. The tissue was rinsed with increasing concentrations of trypsin inhibitor followed by plating medium (Basal Medium Eagle, 5% fetal bovine serum, Glutamax (Invitrogen), N2 supplements (Invitrogen), and penicillin/streptomycin). Cells were plated onto 12mm round German cover glass coated with Matrigel (Corning) at a density of ∼250,000 cells per coverslip. After 4h and every 2-3 days subsequently, 50% of the medium was changed to fresh Neurobasal supplemented with Glutamax, GS21 supplements (AMSBIO), and penicillin/streptomycin.

### Heat shock of worms

NGM plates (8.9ml liquid NGM/plate, weight 13.5±0.2 g) were seeded with 300 µl OP50 in the center and allowed to dry for 48 hrs. Either L4s hermaphrodites were passaged on to these plates or worms were bleach-hatched on to these plates and allowed to grow to Day1 adults. All heat shock experiments were performed with 1-day-old gravid animals. To induce heat shock response in *C. elegans*, NGM plates containing 1-day-old animals were parafilmed and immersed in water bath (product no. ITEMP 4100 H21P 115V, Fischer Scientific, Pittsburgh, PA) pre-warmed to 34°C, for indicated times (5 or 15 min). When required, animals were recovered in 20°C incubators following heat shock, under standard condition after the parafilm was removed. Animals were harvested immediately following heat shock, or following recovery, by rapidly washing them off the plates in sterile water or the appropriate buffer (for ChIP-qPCR and RNA seq experiments) or by picking into 1.5 ml tubes (optogenetics).

### Transfection of mammalian cells

NT2 cells were transfected with Lipofectamine LTX Plus reagent (catalog no. 15338030, Thermo Fisher Scientific) according to manufacturer’s protocol.

### Exogenous 5-HT treatment of worms

As described previously^5^, a 5-HT (catalog no. 85036, Sigma-Aldrich) stock solution of 10mM was made in sterile water, filter-sterilized and then diluted to 2mM before use. This solution (or sterile water as control) was dropped onto the surface of OP50 bacterial lawns (such that the lawns were fully covered in 5-HT) on NGM plates and allowed to dry for ∼2 hrs. at room temperature. Day 1 adult animals were placed onto the 5-HT-soaked OP50 bacterial lawns.

### 5-HT treatment of cells or mouse cortical neurons

Cells were seeded at 1 X 10^5^ cells/ml density the day before the experiment. Cell density influenced the experimental outcome and therefore cell numbers were maintained by counting in the hemocytometer. Since regular serum contains 5-HT, cells were grown in presence of dialyzed fetal bovine serum (Thermo Fisher Scientific) for at least 24 hrs. prior to all experiments. A 5-HT (catalog no. 85036, Sigma-Aldrich) stock solution of 10mM was in sterile water. Cells were incubated at 37°C with different concentrations of 5-HT (or sterile water as control) for different time periods as mentioned in the figures/figure legends and harvested for subsequent assays. For mouse cortical neurons, on the 11^th^ day *in vitro*, each coverslip containing ∼250,000 cells was incubated with different concentrations of 5-HT (or sterile water as control) for different time points. Following this incubation, medium as removed and cultures were immediately harvested for RNA preparation.

### Treatment of mammalian cells with 5-HT agonists

Stock solutions of Sumatriptan succinate (5-HT1 receptor agonist; catalog no. S1198, Sigma-Aldrich), DOI hydrochloride (5-HT2A receptor agonist; catalog no. D101, Sigma-Aldrich), BIMU8 hydrate (5-HT4 receptor agonist; catalog no. B4063, Sigma-Aldrich) and ST1936 (5-HT6 receptor agonist; catalog no. SML0260, Sigma-Aldrich) were made in sterile water or DMSO (also used as control). Cells were grown overnight at described densities in presence of dialyzed fetal bovine serum, incubated with different concentrations of 5-HT agonists for indicated time periods (specified in figures and figure legends) and immediately harvested for experiments. Mouse cortical neurons were treated with BIMU8 hydrate following the protocol for exogenous 5-HT described above.

### Treatment of cells with PKA inhibitor (H89)

Cells were grown overnight in presence of dialyzed fetal bovine serum and then treated with 10 µM H89 (catalog no. B1427, Sigma-Aldrich) for 2hrs. and followed by treatment with 5-HT4 agonist (or control).

### Bleach hatching

*C. elegans* populations contained a large number of gravid adults were selected by picking maintenance plates 5 days after the passage of L4s, as described above. Animals were washed off the plates with 1X PBS and the worms were pelleted by centrifuging at 5000 rpm for 30 seconds. The PBS was removed carefully, and worms were gently vortexed in presence of bleaching solution (250 µl 1N NaOH, 200 µl standard bleach, 550 µl sterile water) until all the worm bodies were dissolved (approximately 5-6 minutes). The eggs were pelleted by centrifugation (5000 rpm for 45 seconds) and bleaching solution was removed. Eggs were washed with sterile water three times and then counted. Care was taken to ensure that all the embryos hatched following this treatment. The eggs were seeded on fresh OP50 or RNAi plates (∼100 eggs/plate for gene expression analysis and ∼ 200 eggs/plate for chromatin immunoprecipitation) and allowed to grow as day-1-adults under standard condition (20°C).

### RNA interference

RNAi experiments were conducted using the standard feeding RNAi method. Bacterial clones expressing the control (empty vector pL4440) construct and the dsRNA targeting different *C. elegans* genes were obtained from the Ahringer RNAi library^6^ now available through Source Bioscience (https://www.sourcebioscience.com/errors?aspxerrorpath=/products/life-science-research/clones/rnai-resources/c-elegans-rnai-collection-ahringer/). *kin-1* RNAi construct was obtained from Dharmacon (catalog no. RCE1182-202302363). All RNAi clones used in experiments were sequenced for verification before use. For RNAi experiments, RNAi bacteria with empty (pL4440 vector as control) or RNAi constructs were grown overnight in LB liquid culture containing ampicillin (100 µg/ml) and tetracycline (12.5 µg/ml) and then induced with IPTG (1 mM) for 2 hours before seeding the bacteria on NGM plates supplemented with ampicillin (100 µg/ml), tetracycline (12.5 µg/ml) and IPTG (1 mM). Bacterial lawns were allowed to grow for 48 hours before the start of the experiment. RNAi-induced knockdown was conducted by (a) dispersing the bleached eggs onto RNAi plates or (b) feeding L4 animals for 24 hours (as they matured from L4s to 1-day-old adults) or (c) feeding animals for over one generation, where second-generation animals were born and raised on RNAi bacterial lawns (*hsf-1*). RNAi-mediated knockdown was confirmed by scoring for known knock-phenotypes of the animals subject to RNAi (slow and arrested larval growth as well as larval arrest at 27°C for *hsf-1* RNAi; dumpy adults for *kin-2* RNAi). *rrf-1(pk1417) I* (NL2098) and *mkcSi13 II; rde-1(mkc36) V* (DCL569) worms were used for germline-specific RNAi experiments whereas *rde-1(ne219) V; kbIs7* (VP303) worms were used for intestine-specific RNAi experiments. These worms were grown in control and *hsf-1* RNAi plates for two generations as mentioned above and day-1 adults were used for heat shock experiments. For germline-specific knockdown of *kin-1* and *kin-2*, *mkcSi13 II; rde-1(mkc36)* worms were bleach hatched on control RNAi or *kin-1*/*kin-2* RNAi plates and experiments were performed with 1-day old animals.

### Knockdown of mammalian HSF1 and SUPT16H by siRNA

Control siRNA and siRNA targeting human *HSF1* and *SUPT16H* (SPT16) were procured from Santa Cruz Biotechnology Inc, USA (catalog no. sc-37007, sc-35611 and sc-37875 respectively) and NT2 cells were transfected with Lipofectamine LTX Plus reagent according to manufacturer’s protocol. All experiments were performed 48 hrs. after transfection and knockdown of endogenous HSF1 and *sp16* was confirmed by western blotting or qRT-PCR respectively. The protein levels were quantified using ImageStudio (LI-COR)

### Optogenetic activation of serotonergic neurons

Optogenetic experiments were performed according to previously published methods as per the requirements of the experiment ^5,7^. Briefly, experimental plates (ATR+) were made from 100 mM ATR (product no. R2500, Sigma-Aldrich) stock dissolved in 100% ethanol and then diluted to a final concentration of 2.5 mM into OP50 or L4440 or *kin-1* or *hmg-3* RNAi bacterial culture and 200µl was seeded onto a fresh NGM plate. Control (ATR-) plates were seeded at the same time with the same culture without adding ATR. All plates were allowed to dry overnight in the dark before use. The *C. elegans* strain AQ2050 was used for this experiment. L4s were harvested on to ATR+ and ATR- plates and the experiment was carried out with day 1 adults. All plates were kept in the dark and animals were allowed to acclimatize to room temperature (20°C to 22 °C) for about 30 min. before starting the experiment. Animals were illuminated with blue light for 30 seconds at a 6.3X magnification using an MZ10 F microscope (Leica) connected to an EL6000 light source (Leica) and harvested at different time points as indicated in Trizol and snap-frozen immediately in liquid nitrogen for RNA extraction. Optogenetic 5-HT release during light stimulation was confirmed by measuring pharyngeal pumping rates.

### Single-molecule fluorescence *in situ* hybridization (smFISH)

smFISH probes were designed against the worm *hsp70* (F44E5.4/5) gene by using the Stellaris FISH Probe Designer (Biosearch Technologies Inc.) available online at http://www.biosearchtech/com/stellarisdesigner. The fixed worms were hybridized with the F44E5.4/5 Stellaris FISH Probe set labelled with Cy5 dye (Biosearch Technologies Inc.) following the manufacturer’s protocol. About 20 wild-type (N2) day 1 worms per condition (control and 34°C heat shock for 5 and 15 min) were harvested by picking off plates immediately after heat exposure into 1X RNase-free phosphate-buffered saline (PBS) (catalog no. AM9624, Ambion), fixed in 4% paraformaldehyde, and subsequently washed in 70% ethanol at 4°C for about 24 hours to permeabilize the animals. Samples were washed using Stellaris Wash Buffer A (catalog no. SMF-WA1-60, Biosearch Technologies Inc.), and then the hybridization solution (catalog no. SMF-HB1-10, Biosearch Technologies Inc.) containing the probes was added. The samples were hybridized at 37°C for 16 hrs, after which they were washed three times with Wash Buffer A and then incubated for 30 min in Wash Buffer A with DAPI. After DAPI staining, worms were washed with Wash Buffer B (catalog no. SMF-WB1-20, Biosearch Technologies Inc.) and mounted on slides in about 16µl of Vectashield mounting medium (catalog no. H-1000, Vector Laboratories). Imaging of slides was performed using a Leica TCS SPE Confocal Microscope (Leica) using a 63X oil objective. LAS AF software (Leica) was used to obtain and view z-stacks.

### Immunofluorescence staining of dissected gonads

Immunostaining of dissected gonads of *C. elegans* was performed to visualize HSf-1::FLAG in the germline. The procedure was conducted as described earlier^5^. Day-1 wild-type adults harboring HSF-1::FLAG and *tph-1* HSF-1::FLAG worms were picked into 15 µl of 1X PBS (pH7.4) on a coverslip, and quickly dissected with a blade (product no. 4-311, Integra Miltex). A charged slide (Superfrost Plus, catalog no. 12-550-15, Thermo Fisher Scientific) was then placed over the coverslip and immediately placed on a pre-chilled freezing block on dry ice for at least 5 min. The coverslip was quickly removed, and the slides were fixed in 100% methanol (−20°C) for 1 min. and then fixed in 4% paraformaldehyde, 1X PBS (pH7.4), 80mM HEPES (pH 7.4), 1.6 mM MgSO_4_ and 0.8 mM EDTA for 30 min. After rinsing in 1X PBST (PBS with Tween 20), slides were blocked for 1 hour in 1X PBST with 1% BSA and then incubated overnight in 1:100 mouse anti-FLAG (catalog no. F1804, Sigma Aldrich) antibody. The next day, slides were washed and then incubated for 2 hrs. in 1:1000 donkey anti-mouse Cy3 (code no. 715-165-150, Jackson ImmunoResearch Laboratories) before they were washed and incubated in DAPI in 1X PBST and then mounted in 10 µl of Vectashield mounting medium (catalog no. H-1000, Vector Laboratories) and imaged as described above.

### Immunofluorescence staining of NT2 cells

NT2 cells grown overnight on coverslips (1 X 10^5^ cells/ml density) in presence of dialyzed fetal bovine serum were fixed in 4% paraformaldehyde in PBS at RT for 10min. Fixed cells were permeabilized with 0.1% Triton-X-100 in PBS at 37°C for 5min, blocked with 1% BSA in PBS at 37°C for 30min, and incubated with rabbit anti-HSF1 antibody (1:100 dilution) (catalog no. 4356, Cell Signaling Technology) for 2hrs. After washing, cells were incubated with AlexaFluor 488-conjugated goat anti-rabbit IgG (H+L) (catalog no. A-11008, Invitrogen) for 2hrs. After washing, coverslips were mounted in Vectashield mounting medium containing DAPI and imaged as mentioned earlier. Images were collected using a Leica Confocal SPE8 microscope using a 63× numerical aperture 1.42 oil-immersion objective lens. The relative intensity of HSF1 in the nuclei of control NT2 cells, and cells treated with BIMU8 in the presence or absence of H89 was quantified from the projections of confocal z-stacks using ImageJ. Background signal was subtracted from each of the projections and the mean intensity for regions corresponding to nuclei of the cells was determined. The average of 25 cells was used to determine the mean intensity for HSF1 staining.

### Assays to evaluate progeny survival following heat shock

#### Progeny survival following 5 minute and 15 minute maternal heat-shock

N2 and *tph-1* L4s were picked on fresh OP50 plates the day before the experiment. After 24-26 hours, 1-day-old animals were either heat shocked at 34°C for 5 or 15 minutes in the water bath or left untreated (control). Heat-shocked animals were either (a) moved to fresh OP50 plates to lay eggs for a 2 hour duration immediately after heat shock (0-2 hr. embryos) or (b) allowed to recover in an incubator at 20°C for 2 hours, and then moved to fresh OP50 plates and allowed to lay eggs for a 2 hour duration (2-4 hrs.). Control embryos were those laid by non-heat shocked animals from the same 2 hour duration. For all experiments except those processed for mRNA, embryos scored were from 5 worms per plate (2 plates per experiment). To score viable embryos, the number of eggs laid were counted, embryos were allowed to hatch at 20°C incubator, and the number of live progeny were scored 48 hrs later. We ascertained that these larvae subsequently grew into adults.

#### Progeny survival following heat-shock following RNAi-induced knockdown in parents, or 5-HT treatment of parents

When RNAi treatment was required, the parents were bleach hatched on fresh RNAi plates and allowed to grow under standard condition (20°C). Animals on Day-1 of adulthood were then transferred onto fresh RNAi plates, subjected to heat-shock or used as controls as described in order to calculate the percentage of live progeny. For assaying recue of *tph-1* embryonic viability by 5-HT treatment, *tph-1* mutant animals were transferred to 5-HT plates made as decrbed above, immediately heat-shocked for 5 minutes at 34°C, allowed to recover for 2 hours on the same plate, and then transferred to fresh OP50 plates without 5-HT to lay eggs for 2 hours. This was because the rate of transit of exogenous 5-HT through the animal is poorly understood.

#### Survival of homozygous and heterozygous progeny

To assess maternal contribution, 5 *tph-1* hermaphrodites (L4s) were allowed to mate with 10 wild-type (N2) males for 26 hrs. Wild-type hermaphrodites (L4) were also allowed to mate with wild-type males or for 26 hrs in similar numbers to control for any effects of mating. Mating was ascertained by counting, post-hoc, the numbers of male progeny laid by these hermaphrodites and ensuring they were ∼ 50% male. The mated hermaphrodites were heat shocked at 34°C for 5 minutes, the males removed, and hermaphrodites allowed to recover for 2 hours at 20°, and then transferred to new OP50 plates to lay eggs for a 2 hour interval. The embryos laid by these hermaphrodites were scored for viability as described above. The hermaphrodites were then transferred to new plates and their male progeny counted so as to ascertain they had indeed mated. Unmated wild-type and *tph-1* hermaphrodites were also heat shocked at the same time. Mated and unmated wild-type and *tph-1* animals that were not subjected to heat shock were used as control.

#### Survival assay to determine the contribution of heat-shocked sperm

Wild-type day-1 males were heat shocked at 34°C for 5 minutes and then transferred onto plates containing L4 wild-type hermaphrodites and allowed to mate for 26 hrs. Mating was ascertained by counting, post-hoc, the numbers of male progeny laid by these hermaphrodites and ensuring they were ∼ 50% male. The gravid 1-day old hermaphrodites were then heat shocked at 34°C for 5 minutes, the males removed, and hermaphrodites transferred immediately onto new OP50 seeded plates to lay eggs for a duration of 2 hours. The hermaphrodites were then transferred to new plates and their male progeny counted so as to ascertain they had indeed mated. Percent viability of embryos was calculated as mentioned earlier.

#### Progeny survival following a prolonged heat exposure

Control and heat-shocked (34°C for 5 minutes), wild-type and *tph-1* day-1 animals were allowed to lay eggs for 2 hours and then all animals were taken off from the plates. After 48 hours, the numbers of progeny that hatched was calculated as described above, and the progeny were then subjected to a prolonged (3 hrs.) heat exposure of 34°C. This condition was chosen after prior experiments to titrate death of control, non-heat shocked progeny to ∼50% to prevent ceiling effects. The percent larvae that survived the prolonged heat shock was scored 24 hours later.

### RNA-sequencing and Data analysis

#### a) RNA isolation, library preparation and sequencing

Age synchronized day 1 adult wild-type, *tph-1(mg280)*II and *hsf-1(sy441)*I animal, upon heat-shock or control conditions, were harvested for RNA extraction. Total RNA was extracted from biological triplicates. Sample lysis was performed using a Tissuelyser and a Trizol-chloroform based method was used in conjunction with the Zymo RNA Clean & Concentrator kit to obtain RNA. The Illumina TruSeq stranded mRNA kit was used to obtain stranded mRNA via Oligo-dT bead capture, and cDNA libraries were prepared from 500ng RNA per sample. Use of stranded cDNA libraries have been shown to maximize the accuracy of transcript expression estimation, and subsequent differential gene expression analysis^8^. Each sample was multiplexed on 6 lanes of the Illumina HiSeq 4000 sequencer, generating 2x150bp paired end reads, with about 43 to 73 million reads per sample.

#### b) RNA-seq analysis

FASTQC was used to evaluate the quality of the sequences. Sequence reads were trimmed of adapters contamination and 20 base pairs from the 5’ and 3’ ends by using Trim Galore Version0.6.0 (www.bioinformatics.babraham.ac.uk/projects/trim_galore/) . Only reads with a quality higher than Q25 were maintained. HISAT2 ^9^ was used to maps the trimmed reads to the C. elegans genome release 35 (WBcel235). On average, 99.4% of the reads mapped to the reference genome. Assemblies of the sequences were done with StringTie ^10^ using the gene annotation from Ensembl WBcel235^11^. DESeq2^12^ was used to identify the genes differentially expressed between the samples. Genes with low read counts (n < 10) were removed from the DESeq2 analysis. Genes with a False Discovery Rate < 0.01 were considered significant. The genes selected for the heatmaps were the genes with significant differences in the wild type control vs wild type heat shock samples. If these genes were not significant in *sy441* control vs *sy441* heat shock or *tph-1* control vs *tph-1* heat shock comparisons, the log_2_foldchange values were adjusted to 0. Principal component analysis (PCA) and pairwise distance analysis (sample-to-sample) were performed by using normalized counts coupled with the variance stabilization transformation (VST). The PCA was done using the top 100 genes with the highest variance in read counts, the pairwise distance analysis was done using the complete set of genes and calculating the Euclidean distance between the replicates.

#### c) Functional analysis

We used the R package clusterProfiler to perform a Gene Ontology (GO) analysis ^13^ on the differentially expressed genes. GO terms with qvalue < 0.05 were considered significant. GO annotations for C. elegans were obtained from R package org.Ce.eg.db: Genome wide annotation for Worm ^14^.

#### d) Data availability

RNA-seq data have been deposited and available at https://dataview.ncbi.nlm.nih.gov/object/PRJNA576016?reviewer=og1kqjgld8qgk0d0qa0ck3di53

### RNA extraction and quantitative real-time PCR (qRT-PCR)

RNA was collected from day-1-adults, and embryos laid by 30-50 animals during 2-4 hours post heat shock. Adult animals were either passaged the previous day as L4s at densities of 20 worms/plate, or were bleach hatched (∼100 eggs/plate). RNA extraction was conducted according to previously published methods ^15^. Briefly, RNA samples were harvested in 50 µl of Trizol (catalog no. 400753, Life Technologies) and snap-frozen immediately in liquid nitrogen. For RNA extraction from embryos, the embryos were subjected to free-thaw cycles five times. The following steps were carried out immediately after snap-freezing or samples were stored at -80°C. Samples were thawed on ice and 200 µl of Trizol was added, followed by brief vortexing at room temperature. Samples were then vortexed at 4°C for at least 45 minutes to lyse the worms completely or lysed using a Precellys 24 homogenizer (Bertin Corp.) according to manufacturer’s protocol. RNA was then purified as detailed in the manufacturer’s protocol with appropriate volumes of reagents modified to 250 µl of Trizol. For RNA extraction from cultured cells and mouse cortical neurons, cells/ neurons were washed with 1X PBS and then harvested in 800 µl of Trizol and snap-frozen in liquid nitrogen. RNA was extracted according to manufacturer’s protocol with appropriate volumes of reagents modified to 800 µl of Trizol. The RNA pellet was dissolved in 17 µl of RNase-free water. The purified RNA was then treated with deoxyribonuclease using the TURBO DNA-free kit (catalog no. AM1907, Life Technologies) as per the manufacturer’s protocol. In case of cultured cells and cortical neurons, 1 µg of total RNA was used for complementary DNA (cDNA) synthesis. cDNA was generated by using the iScript cDNA Synthesis Kit (catalog no. 170-8891, Bio-Rad). qRT-PCR was performed using LightCycler 480 SYBR Green I Master Mix (catalog no. 04887352001, Roche) in LightCycler 480 (Roche) or QuantStudio 3 Real-Time PCR System (Thermo Fisher Scientific) at a 10 µl sample volume, in a 96-well white plate (catalog no. 04729692001, Roche). The relative amounts of *hsp* mRNA were determined using the ΔΔ*C*_t_ method for quantitation. Expression of *pmp-3* (for *C. elegans*) or *GAPDH* (for NT2 cells and mouse primary cortical neurons) was used as internal control. All relative changes of *hsp* mRNA were normalized to either that of the wild-type control or the control for each genotype (specified in figure legends). ΔΔ*C*_t_ values were obtained in triplicate for each sample (technical replicates). Each experiment was then repeated a minimum of three times. For qPCR reactions, the amplification of a single product with no primer dimers was confirmed by melt-curve analysis performed at the end of the reaction. Reverse transcriptase-minus controls were included to exclude any possible genomic DNA amplification. Primers were designed using Roche’s Universal Probe Library Assay Design Center software or Primer3 software and generated by Integrated DNA Technologies. The primers used for the qRT-PCR analysis are listed below:

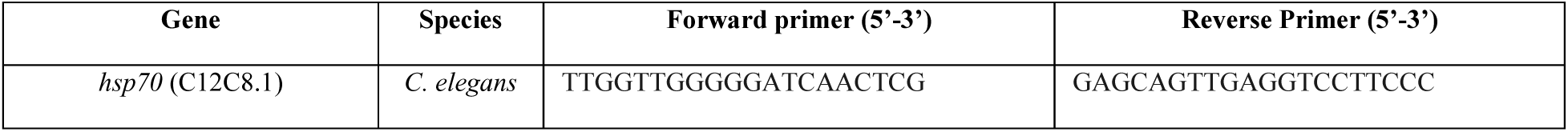
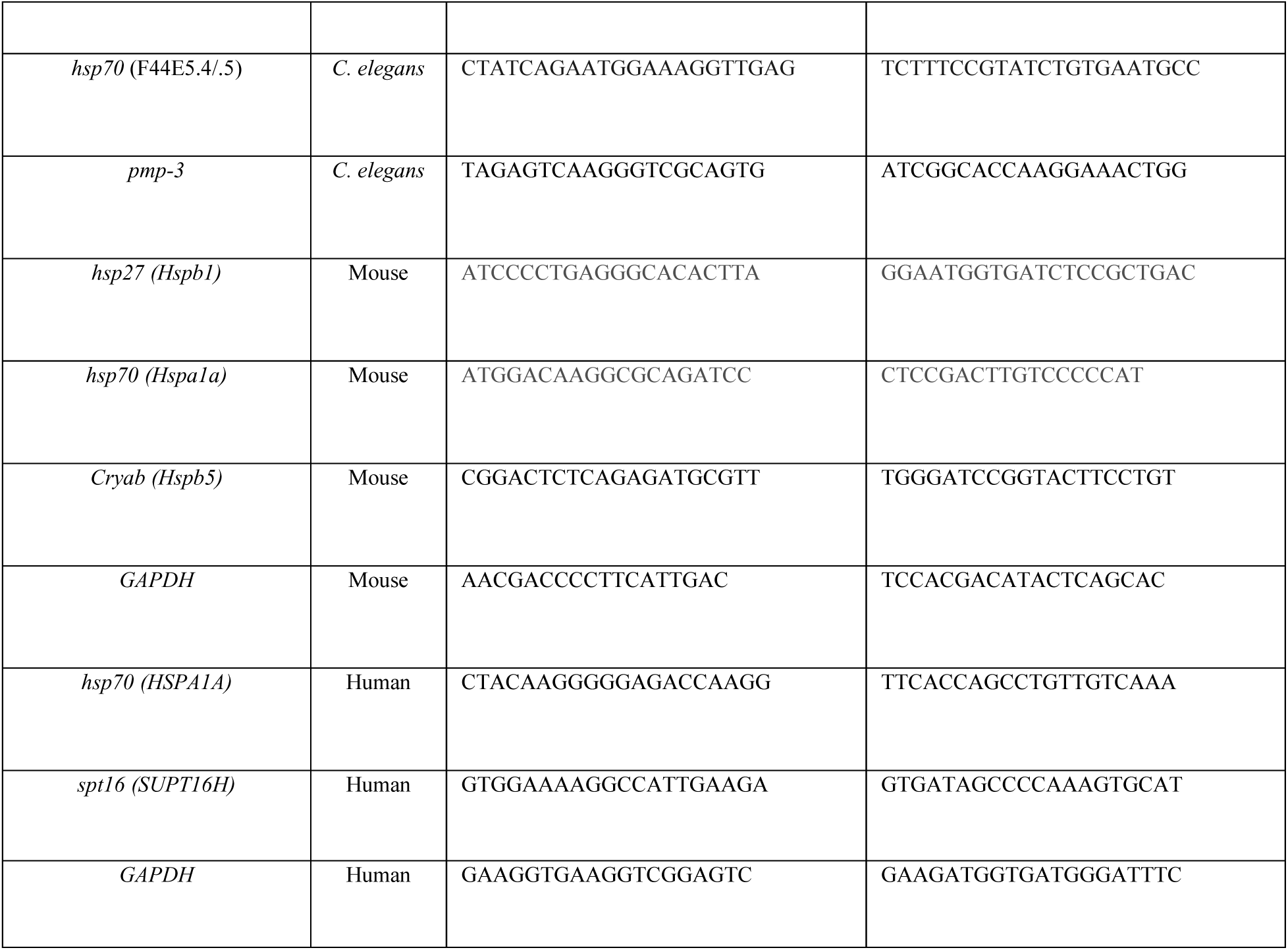

### Western Blotting

Western blot analysis was performed with adult day-1 animals. For protein analysis 20-30 worms were harvested in 15 µl of 1X PBS (pH 7.4), and then 4X Laemmli sample buffer (catalog no. 1610737, Bio-Rad) supplemented with 10% β-mercaptoethanol was added to each sample before boiling for 30 minutes. Whole-worm lysates were resolved on 8% SDS-PAGE gels and transferred onto nitrocellulose membrane (catalog no. 1620115, Bio-Rad). Membranes were blocked with Odyssey Blocking Buffer (part no. 927-50000, LI-COR). Immunoblots were imaged using LI-COR Odyssey Infrared Imaging System (LI-COR Biotechnology, Lincoln, NE). Mouse anti-FLAG M2 antibody (catalog no. F1804, Sigma Aldrich) was used to detect, HSF-1::FLAG. Rabbit anti-HA (catalog no. ab9110, Abcam) was used to detect HMG-3::HA. Mouse anti-α-tubulin primary antibody (AA4.3), developed by C. Walsh, was obtained from the Developmental Studies Hybridoma Bank (DSHB), created by the National Institute of Child Health and Human Development (NICHD) of the National Institute of Health (NIH), and maintained at the Department of Biology, University of Iowa. The following secondary antibodies were used: Sheep anti-mouse IgG (H&L) Antibody IRDye 800CW Conjugated (catalog no. 610-631-002, Rockland Immunochemicals) and Alexa Fluor 680 goat anti-rabbit IgG (H+L) (catalog no. A21109, Molecular Probes, Invitrogen). LI-COR Image Studio software was used to quantify protein levels in different samples, relative to α-tubulin levels. Fold change of protein levels was calculated relative to wild-type/untreated controls.

For western blot analysis of mammalian cells, cells grown (1 X 10^5^ cells/ml density) in presence of dialyzed fetal bovine serum were washed with ice-cold 1X phosphate buffered saline (PBS), scrapped and pelleted by centrifugation at 300g for 3min at 4°C. Cell lysis was carried out using RIPA buffer (50 mM Tris (pH 7.4), 150mM NaCl, 0.1% SDS, 1% NP-40, 0.5% sodium deoxycholate) supplemented with protease inhibitor cocktail (catalog no. 87785, Thermo Fisher Scientific). Protein concentration of whole-cell lysate was measured by Bradford assay (catalog no. 5000006, Bio-Rad) according to manufacturer’s protocol. The preparation of samples, gel run, transfer of proteins to the membrane and imaging was performed as described earlier. Rabbit anti-HSF1 (catalog no. 4356, Cell Signaling Technology) and rabbit anti-HSF1-S320 (catalog no. ab76183, Abcam) were used to detect total and phosphorylated (S320) HSF1 respectively. Mouse anti-Hsp70 antibody (clone 3A3) was a gift from Dr. Richard I Morimoto, Northwestern University. Mouse anti-α-tubulin primary antibody (AA4.3) was used for detection of tubulin which was used as internal control. Fold change of protein levels was calculated relative to wild-type/untreated controls.

### Chromatin immunoprecipitation (ChIP)

Preparation of samples for ChIP was performed by modifying the protocols previously described ^5,16^. Four hundred 1-day-old animals per condition (control or heat shock at 34°C for 5 or 15 minutes) were obtained by washing off two plates of bleach hatched animals, washed with 1X PBS (pH 7.4), and cross-linked with freshly prepared 2% formaldehyde (catalog no. 252549, Sigma Aldrich) at room temperature for 10 min. Reactions were quenched by adding 250mM Tris (pH 7.4) at room temperature for 10 min and then washed three times in ice-cold 1X PBS supplemented with protease inhibitor cocktail and snap-frozen in liquid nitrogen. The worm pellet was resuspended in FA buffer [50 mM HEPES (pH 7.4), 150 mM NaCl, 50mM EDTA, 1% Triton-X-100, 0.5% SDS and 0.1% sodium deoxycholate], supplemented with 1 mM DTT and protease inhibitor cocktail. We discovered during the course of experiments that the presence of a high concentration of EDTA was crucial for consistent yield of DNA, and to prevent the gradual degradation of DNA that otherwise occurred sporadically during the course of the experiments. We attribute this to the presence of resilient DNases that make their way into our preparation due to the culture condition of *C. elegans*. We assessed the quality of the DNA and ChIP with and without these higher concentrations of EDTA to ensure that the concentration of EDTA was not interfering with any other steps of ChIP. The suspended worm pellet was lysed using a Precellys 24 homogenizer (Bertin Corp.), and then sonicated in a Bioruptor Pico Sonication System (catalog no. B0106001, Diagenode) (15 cycles of 30 sec on/off). All HSF-1 ChIP experiments were performed with wild-type (N2) and *tph-1* animals with FLAG tag at the C-terminus of the *hsf-1* gene. Anti-FLAG M2 magnetic bead (catalog no. M-8823, Sigma-Aldrich) was used to immunoprecipitated endogenous HSF1. Beads were first pre-cleared with chromatin isolated from wild-type worms not having any FLAG tag and salmon sperm DNA (catalog no. 15632-011, Invitrogen). Worm lysate was incubated at 4°C overnight with the pre-cleared FLAG beads. For all other ChIP experiments, Protein A/G Magnetic Beads (catalog no. 88802, Pierce) pre-cleared with salmon sperm DNA was used. Pre-cleared lysate was incubated at 4°C overnight with anti-RNA polymerase II (catalog no. 664906, clone 8WG16, Bio legend), anti-Histone H3 (catalog no. ab1791, Abcam), anti-HA (catalog no. ab9110, Abcam) or control mouse (catalog no. sc-2025, Santa Cruz Biotechnology) and rabbit IgG antibody (catalog no. 2729, Cell Signaling Technology) and then pre-cleared magnetic bead was added and incubated for another 3-4 hrs. Beads were washed with low salt, high salt and LiCl wash buffers and then eluted in buffer containing EDTA, SDS and sodium bicarbonate (pH of the elution buffer was adjusted to 11). The elute was incubated with RNase A and then de-crosslinked overnight in presence of Proteinase K. The DNA was purified by ChIP DNA purification kit (catalog no. D5205, Zymo Research). qPCR analysis of DNA was performed as described above using primer sets specific for different regions of *hsp70* (C12C8.1) and *hsp70* (F44E5.4/.5) genes. The primer pair used for amplifying the promoter region of *hsp70* (C12C8.1) gene immunoprecipitated by FLAG beads (for HSF-1 ChIP) was not suitable to amplify DNA immunoprecipitated by RNA polymerase II. Therefore, we used a different primer pair that recognizes slightly downstream region of *hsp70* (C12C8.1) gene for RNA polymerase II ChIP as mentioned in the table below and also in the figure legends. Promoter region of *syp-1* was amplified for all HSF1-ChIP experiments to quantify non-specific binding of HSF-1 (Supplementary Figure 4C). Chromatin immunoprecipitated by all primary antibodies were compared with corresponding rabbit or mouse control IgG to confirm the specificity (Supplementary Figure 9). For all ChIP experiments, 10% of total lysate was used as ‘input’ and chromatin immunoprecipitated by different antibodies were expressed as % input values. All relative changes were normalized to either that of the wild-type control or the control of each genotype (specified in figure legends) and fold changes were calculated by ΔΔ*C*_t_ method. The primers used for ChIP experiments, and the expected amplicon sizes are as follows:

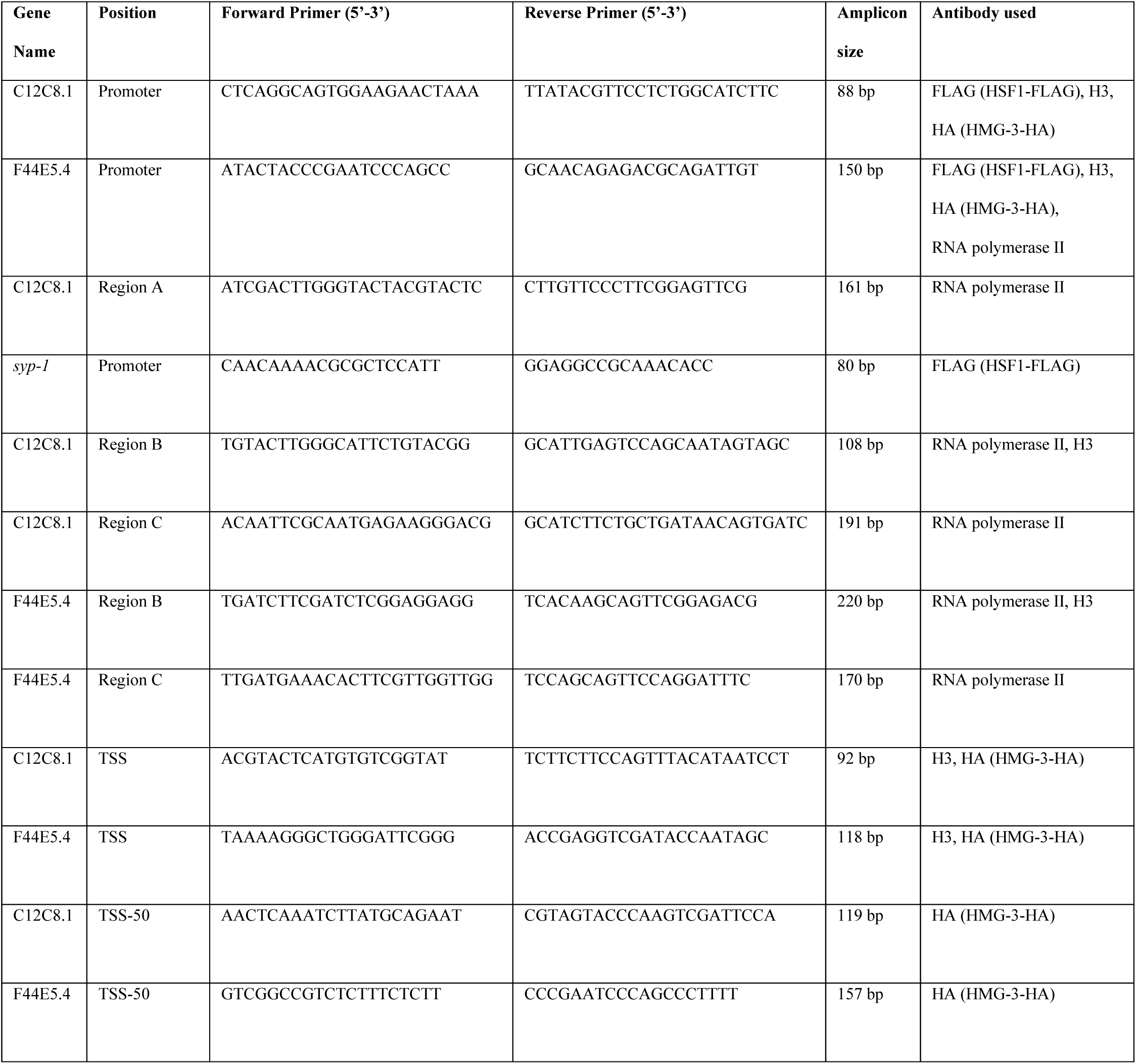

### Statistical analysis

No statistical methods were used to predetermine sample size. The experiments were not randomized. A minimum of three independent experiments (starting from independent parent populations of *C. elegans*) were conducted for all data points. However many experiments were repeated in multiple contexts and n numbers, and mean values reflect all repeats. All qPCR experiments in *C. elegans* were conducted on 30-200 animals per experiment. All ChIP-qPCR experiments were conducted on 200 animals per sample per experiment. For pairwise comparisons such as for qRT-PCR data and progeny hatching, significance was tested using Paired Student’s t tests (assumptions of parametric distributions were first tested and were fulfilled). For all ChIP-qPCR experiments where multiple comparisons were made, significance was tested using one-way ANOVA with Tukey’s multiple comparison correction. Data are indicated as mean ± standard error. p values are indicated as follows: ^∗^p < 0.05, ^∗∗^p < 0.01, ^∗∗∗^p < 0.001. FDR calculations for the RNAseq data set are described in the RNA-seq section of the Methods.

